# Mixed Alkyl/Aryl Phosphonates Identify Metabolic Serine Hydrolases as Antimalarial Targets

**DOI:** 10.1101/2024.01.11.575224

**Authors:** John M. Bennett, Sunil K. Narwal, Stephanie Kabeche, Daniel Abegg, Fiona Hackett, Tomas Yeo, Veronica L. Li, Ryan K. Muir, Franco F. Faucher, Scott Lovell, Michael J. Blackman, Alexander Adibekian, Ellen Yeh, David A. Fidock, Matthew Bogyo

## Abstract

Malaria, caused by *Plasmodium falciparum,* remains a significant health burden. A barrier for developing anti-malarial drugs is the ability of the parasite to rapidly generate resistance. We demonstrated that Salinipostin A (SalA), a natural product, kills parasites by inhibiting multiple lipid metabolizing serine hydrolases, a mechanism with a low propensity for resistance. Given the difficulty of employing natural products as therapeutic agents, we synthesized a library of lipidic mixed alkyl/aryl phosphonates as bioisosteres of SalA. Two constitutional isomers exhibited divergent anti-parasitic potencies which enabled identification of therapeutically relevant targets. We also confirm that this compound kills parasites through a mechanism that is distinct from both SalA and the pan-lipase inhibitor, Orlistat. Like SalA, our compound induces only weak resistance, attributable to mutations in a single protein involved in multidrug resistance. These data suggest that mixed alkyl/aryl phosphonates are a promising, synthetically tractable anti-malarials with a low-propensity to induce resistance.

## Introduction

*Plasmodium falciparum* (*P. falciparum*) is an intracellular parasite responsible for the most severe cases of malaria. While significant progress has been made in reducing malaria cases and fatalities through public health efforts and new treatment strategies over the past two decades, the global malaria burden currently remains on the rise (Rosenthal et al., 2019). In 2021 alone, there were over 247 million reported cases and more than 600,000 deaths (WHO, 2022). Moreover, there has been a continued spread of partial resistance to frontline treatments such as artemisinin (ART) and resistance to other drugs used in artemisinin-based combination therapies (ACTs), underscoring the need to develop agents with novel mechanisms of action that do not induce rapid resistance in the parasite (Ashley et al., 2014; Balikagala et al., 2021; Noedl et al., 2009; Small-Saunders et al., 2022; Straimer et al., 2015).

Natural products have long served as valuable sources for the discovery of potential drugs in *P. falciparum* (Prashar et al., 2022). Salinipostin A (SalA) belongs to a group of bioactive natural compounds known as cyclophostins, characterized by the presence of a bicyclic enolphosphate structure (Spilling, 2019). Notably, these natural products potently inhibit the human serine hydrolase acetylcholinesterase, have antibacterial activity against *mycobacterium*, and have antimalarial properties (Madani et al., 2019; Nguyen et al., 2017; Schulze et al., 2015). We posited that cyclophostins, including SalA, may act via the enolphosphate electrophile to inhibit enzymes. Using an activity-based probe analog of SalA we identified a series of α/β serine hydrolases whose inhibition is responsible for the antimalarial effects of the natural product (Yoo et al., 2020). Furthermore, we found that SalA exhibits a low propensity to induce resistance, a trait shared with artemisinin (ART) and its derivatives. The ability of compounds such as SalA and ART to target multiple proteins likely contributes to the low level of resistance generated (Bridgford et al., 2018; Ismail et al., 2016; Ribbiso et al., 2021). This underscores the potential of targeting multiple serine hydrolases within *P. falciparum* as a potentially valuable therapeutic strategy. However, due to the challenges of isolation and synthesis of SalA, it remains a non-viable lead molecule.

Serine hydrolases are a large superfamily of enzymes that use a catalytic serine to hydrolyze various macromolecules, such as carbohydrates, proteins, and lipids (Long and Cravatt, 2011). In humans, serine hydrolases have been successfully targeted for diabetes and dementia with many candidates for analgesia also in the clinic (Bachovchin and Cravatt, 2012). Nevertheless, serine hydrolases remain underexplored in *P. falciparum*. Serine hydrolases commonly play important roles in lipid metabolism and membrane homeostasis. Throughout the erythrocytic lifecycle of the parasite, the plasma membrane is continuously remodeled to enable the growth of the parasite (Tokumasu et al., 2021). Moreover, *P. falciparum* is unable to synthesize all of its necessary lipids and must scavenge various lipids from the host (Déchamps et al., 2010b, 2010a). Therefore, the parasite likely dynamically regulates the expression and activity of metabolically important enzymes at various points in the lifecycle to ensure proper membrane composition and lipid homeostasis. The activity of serine hydrolases in parasites was first investigated with a broad spectrum fluorophosphonate-desthiobiotin probe which enabled the identification of over 20 active serine hydrolases in the lysate of late-stage schizont parasites (Elahi et al., 2019). More recently, a fluorphosphonate probe was used to identify serine hydrolases in live parasites at different stages within the life cycle, clearly demonstrating the dynamic regulation of lipolytic serine hydrolases in *P. falciparum* (Davison et al., 2022).

A wide range of strategies and warheads have been employed for the development of covalent inhibitors targeting serine hydrolases (Faucher et al., 2020). Phosphonates are a valuable bioisostere because they can mimic phosphate groups, while maintaining reactivity with catalytic serine residues (Elliott et al., 2012). In fact a range of diphenyl phosphonates have been developed to inhibit serine proteases (Jackson et al., 1998; Senten et al., 2003; Serim et al., 2013). Therefore, we postulated that phosphonates could be used as a replacement for the enolphosphate group of SalA (Figure 1A). However, only the broadly reactive fluorophosphonate has been used for activity-based protein profiling (ABPP) and the characterization of serine hydrolases within *Plasmodium* (Lu et al., 2021). The mixed alkyl/aryl phosphonates are a relatively recently reported class of phosphonate-based serine hydrolase inhibitors. This class of serine-specific electrophiles has great potential for use in serine hydrolase inhibitors because reactivity can be tuned through the substitution of various phenol leaving groups appended to the phosphonate (C. Wang et al., 2019). Similar to the widely used carbamates and ureas, the leaving group of the electrophile can be optimized to tune both the selectivity and potency of inhibitors toward specific serine hydrolases (Adibekian et al., 2011; Cognetta et al., 2015).

**Figure 1:**
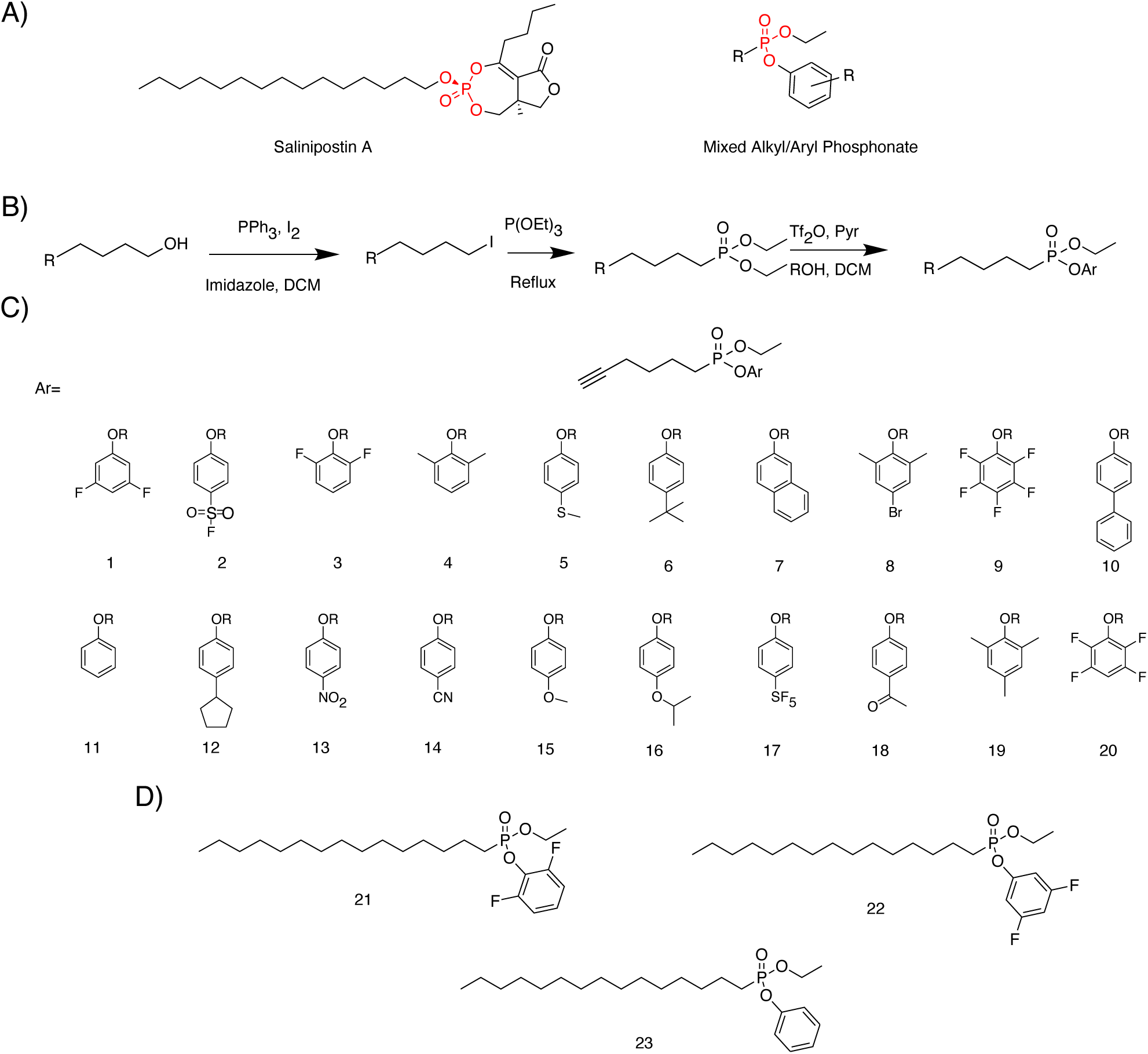
Structure of Salinipostin A and Synthesized Phosphonates. (A) Structure of Salinipostin A and General Mixed Alkyl/Aryl Phosphonate. Salinipostin A (left) is drawn with electrophilic enolphosphate colored red and the mixed alkyl/aryl phosphonate (right) with electrophilic phosphonate group in red. (B) Synthesis of mixed alkyl/aryl phosphonates and structures of molecules. PPh_3_, triphenyl phosphine; I_2_, iodine; DCM, dichloromethane; P(OEt)_3_, triethyl phosphite; Tf_2_O, triflate anhydride; pyr., pyridine; ROH, phenol. (C) Structures of Short Chain Hexyne-based Phosphonates (D)Structures of Pentadecane-based Phosphonates

We thus set out to develop mixed alkyl/aryl phosphonates that could recapitulate the antimalarial properties of SalA but that would be more synthetically tractable. We used various phenol leaving groups and different alkyl chain lengths to investigate the structure-activity relationship of our molecules on purified enzymes and their phenotypic effects on *P. falciparum* parasites. Interestingly, we identified a pair of mixed aryl/alkyl phosphonates that are constitutional isomers with similar biochemical potency against purified serine hydrolases but dramatically different potencies when used on intact parasites. Using these compounds, we performed chemoproteomics to identify a serine hydrolase, abH112, as the primary anti-malarial target of the compound with potent anti-malarial activity. Interestingly, conditional knock down of abH112 did not result in growth defects suggesting that the killing effect of our compound is likely mediated by inhibition of multiple targets which must include abH112. Phenotypic analysis and metabolic profiling studies demonstrated that our mixed phosphonate compound kills parasites through a mechanism that is distinct from that of SalA or the pan-lipase inhibitor Orlistat. Drug pressure selections demonstrated that the antimalarial mixed phosphonate generated resistant parasites with only minor shifts in sensitivity to the compound that were mediated by the same drug resistance protein identified in SalA resistant parasites. This result is consistent with a mechanism of killing involving multiple targets and provides further support for targeting serine hydrolases for anti-malarial therapeutics.

## Results

### Short Chain Mixed Alkyl/Aryl Phosphonates are potent inhibitors of PfMAGL

While synthesis routes have been reported for Salinipostins and other similar natural products, synthesis of the bicyclic enolphosphate group remains the most significant challenge of the total synthesis (Zhao et al., 2018). Additionally, limited product yield makes salinipostins unlikely to be used as therapeutics. We therefore developed a synthetic strategy for the rapid production of mixed alkyl/aryl phosphonates that mimic the main core functional elements of SalA (Figure 1B). This route utilizes an Appel reaction to convert the terminal hydroxyl of an alkyl chain, into an iodine which can then be converted to the corresponding diethyl phosphonate using an Arbuzov reaction. Subsequent aryloxylation using triflate anhydride and pyridine enable us to develop a library of racemic mixed alkyl/aryl phosphonates (Figure 1C; Huang et al., 2018). Our initial library comprised 20 hexyne-based mixed phosphonates with phenolic leaving groups varying in their stereoelectronics.

To characterize the compounds, we conducted a biochemical screen against PfMAGL, a serine hydrolase identified as a primary target of SalA. PfMAGL is annotated as an essential protein based on a piggyBac transposon saturation screen of essential genes in *P. falciparum*, and we previously showed that the biochemical inhibition of PfMAGL correlates with potency against intraerythrocytic parasites (Yoo et al., 2020; Zhang et al., 2018). We screened our library at high (25 µM) and medium (2.5 µM) concentrations and observed that phosphonates with electron-withdrawing substitutions on the phenols effectively inhibited the enzyme (with the exception of compound **13**) whereas those with either electron donating or sterically bulky groups did not (Figure S1A). Compound **2**, containing a p-sulfonyl fluoride substitution, had an IC_50_ of 20 nM and was the most potent inhibitor of PfMAGL in our initial set of compounds (Figure S1B). We obtained similar results screening against the human homolog, hMAGL, with only the phosphonates bearing an electron-withdrawing group exhibiting inhibitory effects and compound **2** showing the highest potency against the human enzyme (Figure S1C; S1D).

Following biochemical characterization of our compounds, we assessed their effects against blood stage parasites in a 72-hour parasite growth assay (Figure S1E). Only compound **2** had significant antiparasitic effects, killing over 50% of the parasites at the 10 µM concentration. In contrast, most compounds, including those with nanomolar inhibition of PfMAGL, showed limited to no killing at both 10 µM and 1 µM, suggesting that inhibition of PfMAGL may not be sufficient for parasite killing. Compound **2** exhibited moderate potency as an inhibitor of parasite growth, with an EC_50_ of 1.6 µM (Figure S1F). However, this compound is problematic due to the fact that it contains an electrophilic p-sulfonyl fluoride that can react with nucleophilic amino acid residues other than the catalytic serine of the intended target (Lou and Willis, 2022; Narayanan and Jones, 2015). Thus, some of the potent anti-parasitic activity may be the result of off target inhibition or general toxicity from the reactive p-sulfonyl fluoride group.

### Long chain lipids containing the mixed phosphonate electrophile are potent inhibitors of PfMAGL and parasite growth

Since our initial set of molecules did not exhibit potent antiparasitic properties, we synthesized a second set of mixed alkyl/aryl phosphonates with a longer alkyl chain on the phosphorus to better mimic the structure of SalA. We chose a C15 lipid to match the length of the western chain on SalA. The synthesis of long chain phosphonates proved to be more challenging, but we were able to synthesize three C15 mixed phosphonates (Figure 1D; Figure 2A). We first tested these compounds for direct inhibition of recombinant PfMAGL. Two constitutional isomers, **21**, with a 2,6-difluorophenol, and **22**, with a 3,5-difluorophenol, were picomolar inhibitors of the enzyme (Figure 2B). Compound **23**, with a phenol leaving group, had only low micromolar potency, and its short-chain counterpart 11 showed no inhibition at concentrations up to 25 µM (Figure S2A), suggesting that the combination of warhead and lipid chain length contribute to the potency of the compounds. We tested the three compounds for inhibition of hMAGL and found similar trends in potency (Figure S2B).

**Figure 2:**
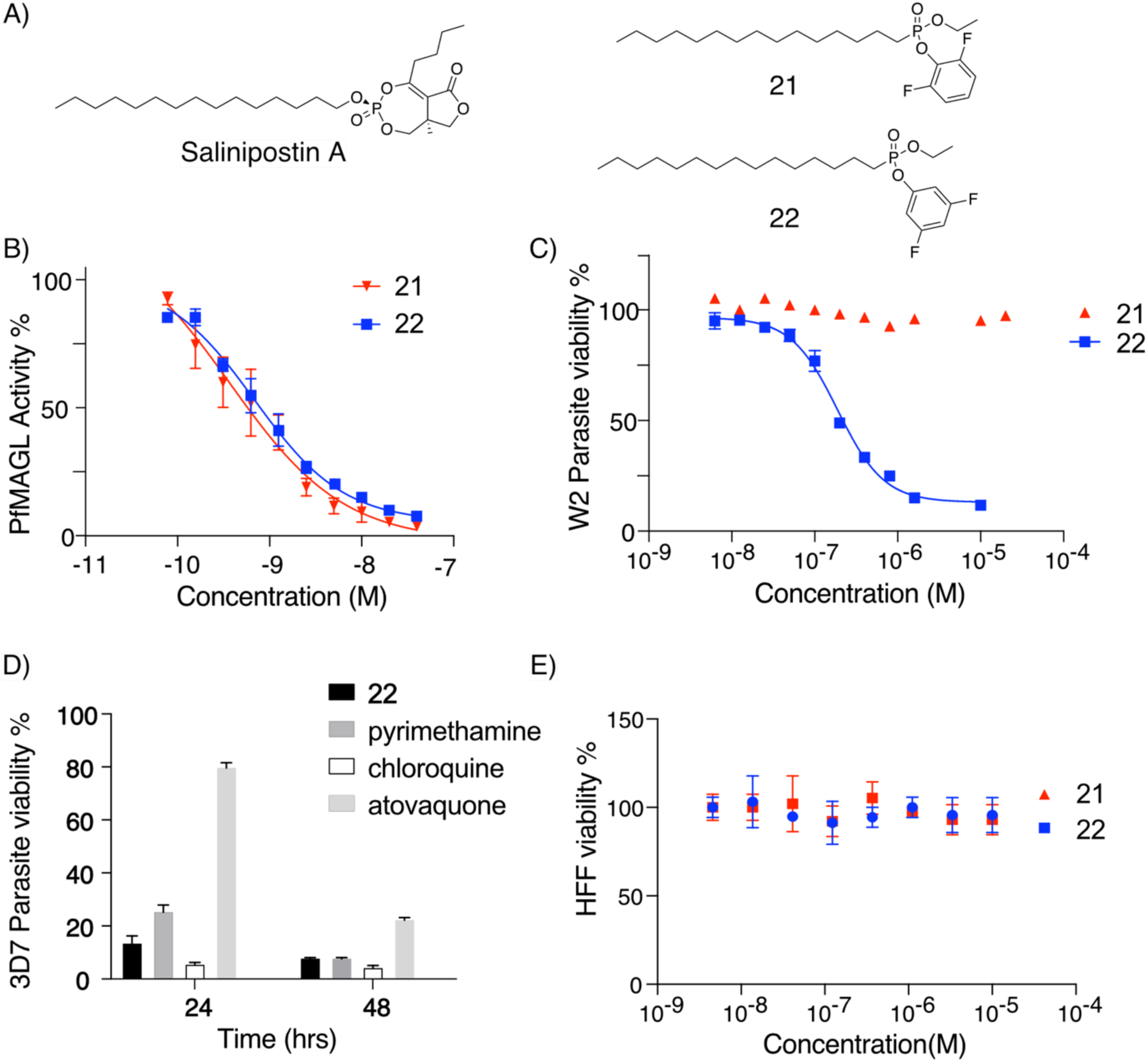
3,5 difluorophenol Substituted Long Chain Phosphonate is a potent inhibitor of parasite growth. (A) Strucutre of Salinipostin A and constitutional isomers of long chain mixed alkyl/aryl phosphonates. (B) IC_50_ curves of short chain mixed alkyl/aryl phosphonates for inhibition of 4-methylumbelliferyl caprylate processing by recombinant PfMAGL. (mean ± SD, N,n = 2,3). (C) EC50 values of Compound 21 and 22 in 72 in a 72 hour treatment of synchronized ring stage W2 parasites with error bars, SEM. (mean ± SD, n=4 from 3 independent experiments). (D) Rate of Kill of Compound 22 measured against compared against known fast (chloroquine), medium (pyrimethamine), and slow (atovaquone) acting inhibitors of parasite growth in which compounds were dosed at their EC_50_ and parasite viability was monitored over time and ± SEM. N,*n* = 6,2. (E) HFF Toxicity for alkyl cyclic peptide inhibitor and linear counterparts. Points are plotted as mean ± SEM values of each inhibitor for HFF growth. Data were generated with 72 hour assays (N, n = 2,3).

After characterizing the biochemical activities of our compounds, we tested the long chain phosphonates in the 72-hour parasite growth assay. Surprisingly, we observed that compound **22** was an extremely potent inhibitor of parasite growth (EC_50_ = 188 nM), while the constitutional isomer **21**, showed no killing at concentrations up to 20 µM (Figure 2C). These results further suggest that even though PfMAGL may be an essential protein, inhibition of this enzyme alone does not seem to be sufficient to kill parasites. In addition, we reasoned that these two isomers would be ideal tools for use in comparative chemoproteomic experiments to identify the protein target(s) responsible for the observed anti-parasitic activity. In fact, this strategy of using a paired set of isomers of an electrophilic small molecule is has recently been demonstrated to be an effective strategy to find phenotypically relevant targets when compounds react with multiple proteins (Y. Wang et al., 2019).

To further investigate the mechanism of the antiparasitic activity of **22,** we used a rate of kill assay. These results confirmed that **22** kills parasites at a rate that is faster than pyrimethamine, but not as fast as chloroquine, suggesting that our inhibitor has the potential to quickly clear an infection (Figure 2D; Sanz et al., 2012). Moreover, when parasites were treated with compound **22** for 1 hour followed by a washout and then continued growth for 72 hours, we observed only a ten-fold drop in potency (EC_50_ = 1.33 µM) compared to continuous treatment, suggesting rapid and prolonged binding to targets consistent with a covalent mechanism of action (Figure S2C). Lastly, we found that, at concentrations up to 50 µM, we saw no impact on mammalian cell viability for either compound **22** or **23**. This result is consistent with studies of SalA and other human serine hydrolase inhibitors, which have no cytotoxic effects on human cells.

### Activity-Based Probes Identify Differentially Targeted Serine Hydrolases

To perform a chemoproteomic identification of the targets of our mixed phosphonates, we synthesized terminal alkyne bearing derivatives that would enable click chemistry to isolate labeled targets. Using a synthetic route based on the previous synthesis of light activated probes of ceramides (Supplemental Scheme 1), we synthesized activity-based probe versions of compounds **21** and **22** (**21-alk** and **22-alk**; Figure 3A; Haberkant et al., 2016). Both compounds had similar potencies against the purified PfMAGL and hMAGL compared to the original non-alkynlyated inhibitors suggesting that addition of the tag did not impact target binding (Figure S3A). We also confirmed that probe **21-alk** remained inactive in the parasite killing assay, and compound **22-alk** remained active with only a modest loss in potency (EC_50_ = 1.04 µM) compared to the parent compound (Figure S3B). This change in potency upon addition of the alkyne resulted in a decrease in potency of a magnitude that was similar to what was observed when an alkyne tag was added to the Salinipostins (Schulze et al., 2015). Regardless of the drop in potency, compound **22-alk** remained a potent inhibitor of *P. falicparum* growth and retained the fast rate of kill observed for the parent compound (Figure S3C).

**Figure 3:**
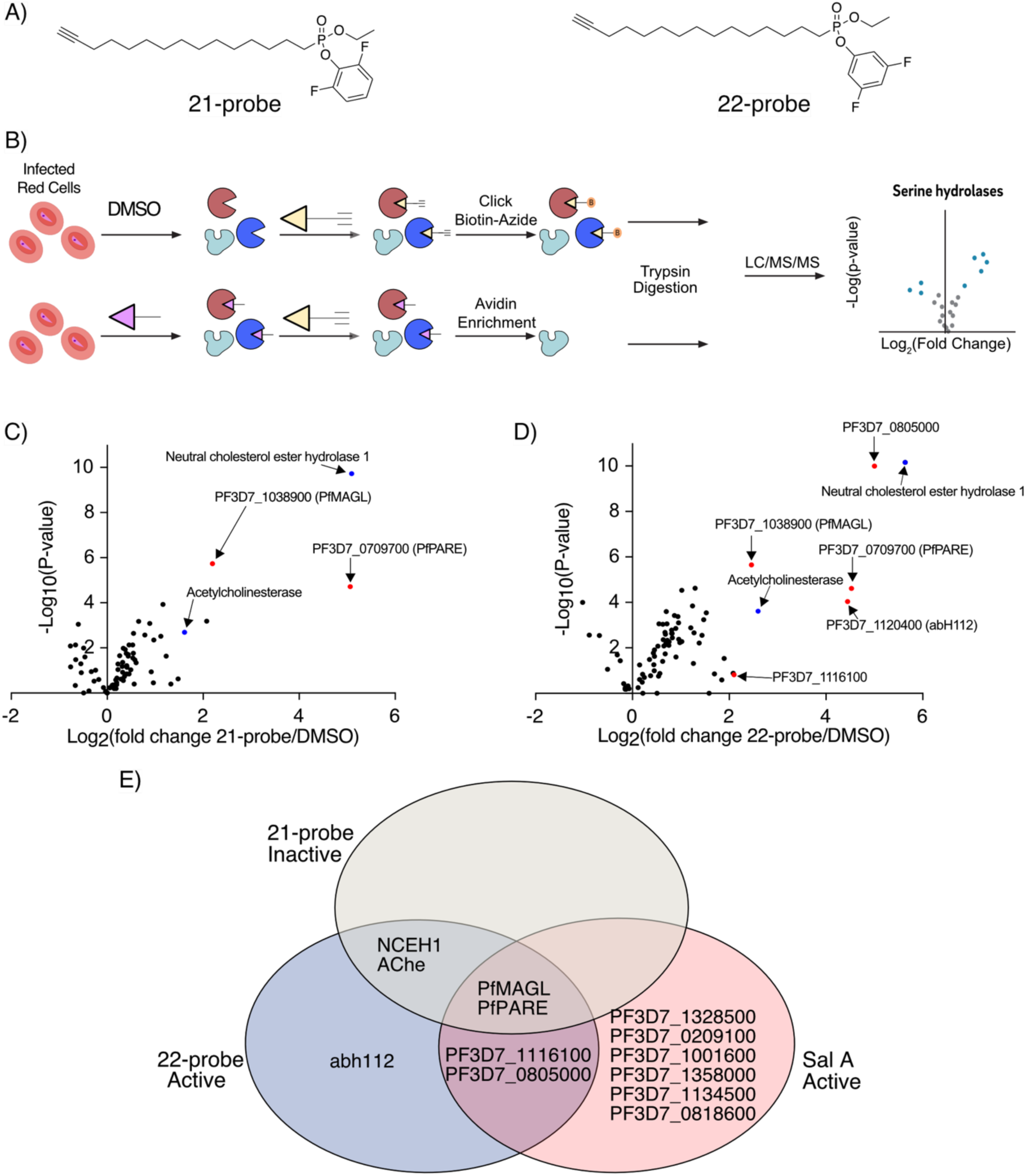
Chemoproteomics identifies distinct serine hydrolases between active and inactive difluorophenol isomers. (A) Structures of Activity-Based Probes used for chemoproteomics. (B) Workflow for Competitive ABPP experiment. Infected red bloods are either pretreated with DMSO or the active or inactive inhibitor (green) and followed by treatment with an activity-based probe (orange), before being submitted through typical proteomics workflows. (C) Volcano plots of Activity-based probe **21-alk** targets in *P. falciparum*. The x axis shows the logarithm values of between **21-alk**-treated and DMSO vehicle-treated parasites. The statistical significance was determined without correcting for multiple comparisons in triplicates and the y value in the volcano plot is the negative logarithm of the p value. Significantly enriched serine hydrolases are denoted in red for parasite enzymes and blue for human enzymes. (D) Volcano plots of Activity-based probe **22-alk** targets in *P. falciparum*. The x axis shows the logarithm values of between **22-alk**-treated and DMSO vehicle-treated parasites. The statistical significance was determined without correcting for multiple comparisons in triplicates and the y value in the volcano plot is the negative logarithm of the p value. Significantly enriched serine hydrolases are denoted in red for parasite enzymes and blue for human enzymes. (E) Venn Diagram showing overlap of significantly enriched serine hydrolases between the inactive probe **21-alk** red, the active probe **22-alk**, and the natural product Salinipostin A.

To identify the target(s) responsible for the anti-parasitic effects of **22**, we performed a competitive ABPP proteomic experiment in infected red blood cells by pretreating with either vehicle or **21** or **22**, before labelling with **21-alk** or **22-alk** (Figure 3B). When labeling with the activity-based probe **22-alk**, we identified seven putative serine hydrolases and one uncharacterized protein (PF3D7_0619300) as being significantly enriched (Figure 3C). The biologically inactive probe **21-alk**, significantly enriched four of those serine hydrolases (Figure 3D). Both probes enriched two human serine hydrolases, Neutral cholesterol ester hydrolase 1 and acetylcholine esterase both of which were previously identified in ABPP experiments and whose inhibition has little effect on parasite growth (Davison et al., 2022). Moreover, both probes labeled PfMAGL (PF3D7_1038900) and the lysophospholipase PfPARE (PF3D7_0709700). PfPARE was previously identified as an esterase that activates the antimalarial drug pepstatin (Istvan et al., 2017). However, its esterase activity is dispensable for parasite growth. This finding coupled with the fact that it was targeted by both the active and inactive probes suggest that it is likely not a relevant target for the observed anti-parasitic activity of our compounds (Figure 3E). PfMAGL, on the other hand, has been previously identified as an essential protein in a PiggyBac transposon saturation screen, and we demonstrated that the biochemical inhibition of PfMAGL is associated with the inhibition of parasite growth when other serine hydrolases are also inhibited (Yoo et al., 2020; Zhang et al., 2018). Interestingly, labeling of PfMAGL was competed by the inactive compound **21** while active inhibitor **22** did not compete, suggesting **22** has stronger affinity for other serine hydrolases (Figure S3 E,F). This suggests that inhibition of PfMAGL alone is not sufficient for parasite killing. Two serine hydrolases (PF3D7_0805000 and PF3D7_1116100) enriched by **22-alk** were also identified as targets of SalA but neither were competed by **22** nor were they annotated as essential in the PiggyBac screen (Figure 3E; Table 1; Yoo et al., 2020). Of the proteins competed by **22**, only PF3D7_1120400, previously identified as AbH112, was annotated as essential, was competed only by the active inhibitor **22**, and was not identified as a target of SalA (Table 1; Figure S3G). Therefore, our proteomic data suggests that abH112 is likely the primary target of **22** that, together with other essential serine hydrolase targets such as PfMAGL, mediates the antiparasitic activity of this mixed akyl/aryl phosphonate compound.

**Table 1.**
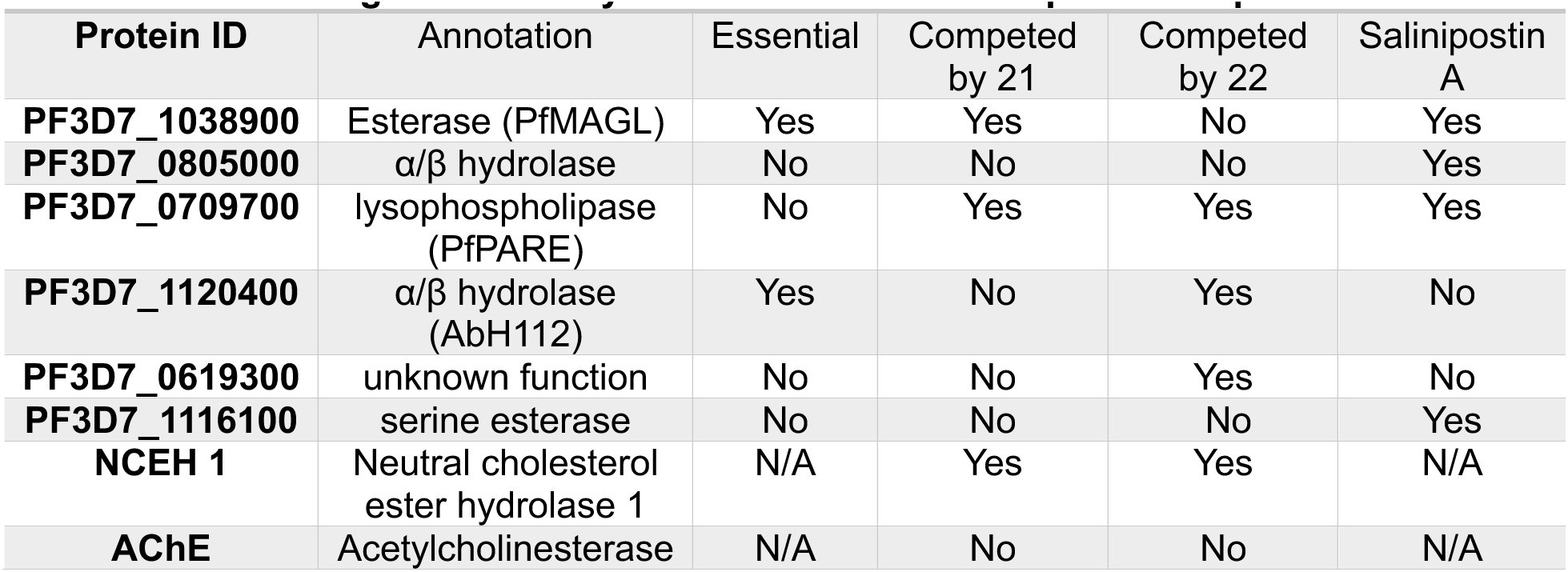
Targets of Activity-Based Probes and Competition Experiments.

| **Protein ID** | **Annotation** | **Essential** | **Competed by 21** | **Competed by 22** | **Salinipostin A** |
| --- | --- | --- | --- | --- | --- |
| **PF3D7_1038900** | Esterase (PfMAGL) | Yes | Yes | No | Yes |
| **PF3D7_0805000** | $\alpha/\beta$ hydrolase | No | No | No | Yes |
| **PF3D7_0709700** | lysophospholipase (PfPARE) | No | Yes | Yes | Yes |
| **PF3D7_1120400** | $\alpha/\beta$ hydrolase (AbH112) | Yes | No | Yes | No |
| **PF3D7_0619300** | unknown function | No | No | Yes | No |
| **PF3D7_1116100** | serine esterase | No | No | No | Yes |
| **NCEH 1** | Neutral cholesterol ester hydrolase 1 | N/A | Yes | Yes | N/A |
| **AChE** | Acetylcholinesterase | N/A | No | No | N/A |

### AbH112 is predicted to be an essential gene but knock down of the protein does not impact parasite growth

To gain insight into the possible function of abH112, we performed bioinformatic analysis of the enzyme. We compiled a list of all the proteins in human, *P. falciparum*, and the related *Toxoplasma gondii* proteomes that share the Protein family (PFam) domain of Hydrolase 4 (PF12146) with AbH112 (Mistry et al., 2021). Using the Clustal Omega multiple sequence alignment algorithm, we performed phylogenetic analysis of the putative serine hydrolase protein sequences. (Figure 4A; Figure S4A). The two proteins with the most homology to AbH112 are the uncharacterized *Taxoplasma gondii* protein TGVAND_223510 and the human serine hydrolase ABHD17A. The human ABHD17A is of particular interest because it was recently identified as a depalmitoylase for nRAS and we have previously shown that depalmitoylases are viable drug targets in *Toxoplasma gondii* (Child et al., 2013; Onguka et al., 2021; Remsberg et al., 2021). However, no depalmitoylases have been identified in *P. falciparum*. We overlayed the AlphaFold predicted structures for both ABHD17A and AbH112 and found that the catalytic triad aspartate, histidine, and serine are aligned (Figure 4B)(Jumper et al., 2021). Furthermore, both proteins show good alignment of lipid binding motifs, the catalytic triad, as well as the GxSxG serine hydrolase motif (Figure 4C). It should be noted that AbH112 contains a GHSxG serine hydrolase motif which has been found in enzymes with dual serine hydrolase and acetyl transferase activity, suggesting a possible function beyond hydrolysis (Poust et al., 2014). To assess whether AbH112 has a function in regulating global palmitoylation, we incubated infected red blood cells for 16 hours with 17-ODYA to incorporate the clickable analog of palmitate into the proteome. We then treated the parasites with either vehicle, compound **21**, **22**, or the palmitoyltransferase inhibitor 2-bromopalmitate (2-BP) before lysing and clicking on TAMRA-azide for a fluorescent SDS-PAGE readout. While we observed changes in patterns of palmitoylation upon treatment with 2-BP, we did not observe any changes in global palmitoylation between the vehicle and the two inhibitor treated samples, suggesting that the inhibition of AbH112 by compound **22** likely does not impact global palmitoylation (Figure S4C).

**Figure 4:**
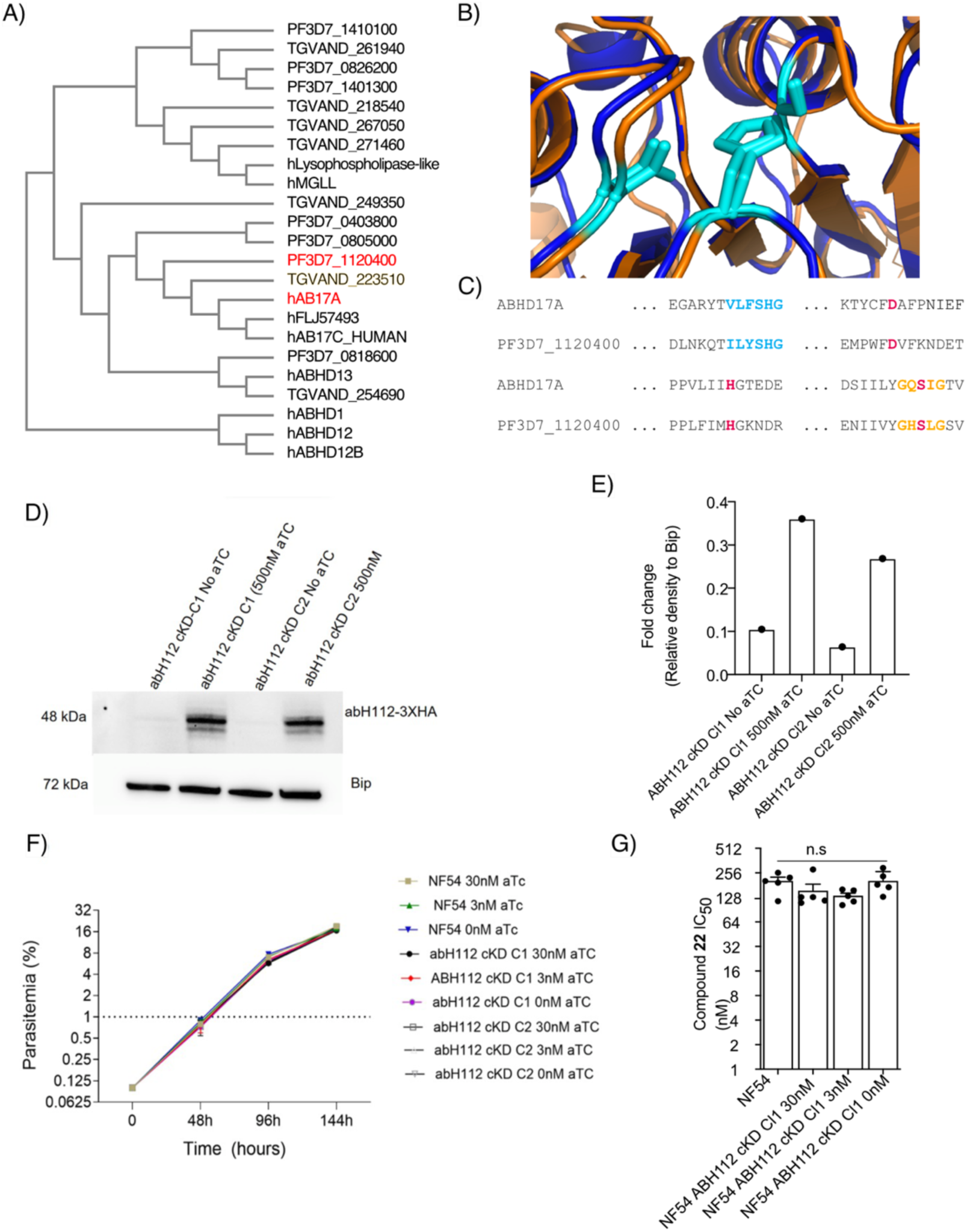
**Conditional Catalytic Knockout of abH112 leads to Gene Duplication.** (A) Partial Dendrogram of *P. falciparum*, *Toxoplasma gondii*, and human serine hydrolases that share a protein family domain generated using the Clustal Omega algorithm with abH112 and hAB17A colored in red. (B) Overlay of predicted AlphaFold structure of abH112 (orange) and hAB17A (blue) with active site catalytic triad in cyan. (C) Sequence alignment of abH112 and hAB17A. Lipid binding motif is colored blue, GXSXG serine hydrolase motif is shown in gold, and catalytic triad is shown in red. (D) Western blotting of abH112 cKD lines grown in +/-aTC. Data were measured by ChemiDoc^TM^ MP Imaging System (Biorad), with bip used as a loading control. (E) Plot of abH112 expression levels from (D) normalized to Bip. (F) Plot of growth levels of parent WT cells (NF54) and abH112 cKD parasites in the presence or absence of aTC. Means are plotted ± SEM (n = 3). Tests for significance used a two-way-ANOVA. G) Plot of EC_50_ values of parasites for compound **22** in the WT parent cells (NF54) and in the cKD lines in the presence and absence of aTc. EC_50_s are plotted as means ± SEMs (N, n = 4-5, 2) and statistical significance determined using Mann-Whitney *U* tests comparing the parental data vs the abH112 cKD line where \**p* < 0.05.

To further characterize abH112, we attempted to express and purify the recombinant enzyme, but we were unable to isolate active forms of the protein. A recent study suggested that the catalytic activity of abH112 may be dispensable to the survival of the parasite (Davison et al., 2022). This study made use of a conditional mutant line in which a loxP DiCre system was used to conditionally replace the catalytic serine-227 with an alanine to catalytically inactivate abH112 (Matera and Wang, 2014; Pieperhoff et al., 2015). This report, however, did not confirm that the abH112 catalytic activity was disabled after recombination nor whether the wild-type gene had been fully excised from the parasite population. To explore this further, we propagated two parasite loxP-tagged recombinant clones (kindly provided by Dr. M. Blackman, the Francis Crick Institute, London) in the presence of rapamycin treatment, which causes recombination between the loxP sites (inserted in artificial introns) and leads to introduction of the mutated coding sequence fragment containing the mutated alanine (Figure S4D). PCR analyses of these parasites demonstrated the correct length for the excised product in rapamycin-treated parasites (Figure S4E). However, we also observed PCR products corresponding to the wild-type sequence in both pre and post-excision samples, which suggests an incomplete recombination or duplication of the gene (Figure S4F).

To further investigate the role of abH112 in asexual blood stage parasites, we used the TetR-DOZI aptamer system to generate cell lines in which the level of expression of the gene is conditionally knocked down (cKD) (Figure S5A; Ganesan et al., 2016). We confirmed site-specific integration by diagnostic PCR (Figure S5B and Supplementary table 1). The TetR-DOZI aptamer system regulates the expression of the target gene through anhydrotetracycline (aTC). Withdrawal of aTC from the culture drastically reduced the level of abH112 protein expression as determined by western blot. However, this reduction in expression of abH112 did not result in any developmental delay or growth defect in asexual blood stage parasites compared to control (Figure 4D-F). This lack of a growth phenotype in abH112 knockdown parasites may be the result of residual abH112 activity or functional redundancy in serine hydrolases during the erythrocytic cycle. To test whether the conditional knockdown impacted global serine hydrolase activity, we performed fluorescent ABPP on lysates of the knockdown parasites under both induced and uninduced expression of abH112. Using the broad-spectrum fluorophosphonate probe, we observed that there were no differences in global serine hydrolase activity between the induced and uninduced expression of abH112, suggesting that knockdown of abH112 does not lead to compensatory increase in the activity of other serine hydrolases. We also tested the susceptibility of abH112 cKD and control NF54 parasites to compound **22** in the presence and absence of aTC. We found that reduction in the levels of abH112 protein did not affect parasite susceptibility to compound **22**, suggesting that this compound likely acts by inhibiting abH112 in combination with other essential serine hydrolase(s) (Figure 4G).

### The Mechanism of Parasites killing by compound 22 is Distinct from the Pan-Lipase Inhibitor Orlistat

Because abH112 is a target that was not identified as a target of SalA, we reasoned that its mode of action is likely distinct. Therefore, we performed a detailed phenotypic analysis of infected red blood cells treated with either inactive compound **21** or active inhibitor **22.** We visualized Giemsa-stained blood smears every twelve hours to determine the blood stage where growth was arrested. Ring stage parasites treated with the inactive **21** progressed through the blood stage lifecycle, egressed, and were able to successfully reinvade new red blood cells (Figure 5A). In contrast, parasites treated with **22** were arrested in growth at 24 hours, corresponding to a block in the transition from the late ring to early trophozoite stage. Previous studies have shown that depletion of palmitate and oleate lead to similar phenotypes (Mi-Ichi et al., 2007). However, supplementation of both fatty acids did not rescue the killing phenotype of **22** (Figure S6A). A more recent report showed that the loss of the *P. falciparum* lysophospholipase, PfLPL1, caused parasite growth arrest in late ring stage via inhibition of food vacuole formation (Asad et al., 2021). Therefore, we evaluated the food vacuoles of parasites treated with either the vehicle, our inactive compound, or our active compound. We observed a small reduction in food vacuole size only in parasites treated with **22**. However, this reduction was not significant and is likely not the primary mechanism associated with parasite killing (Figure S6B-C). Our results suggest a distinct mode of action of killing by **22** compared to both SalA and the pan-lipase inhibitor, Orlistat, both of which block progression at the late schizont stage (Yoo et al., 2020).

**Figure 5:**
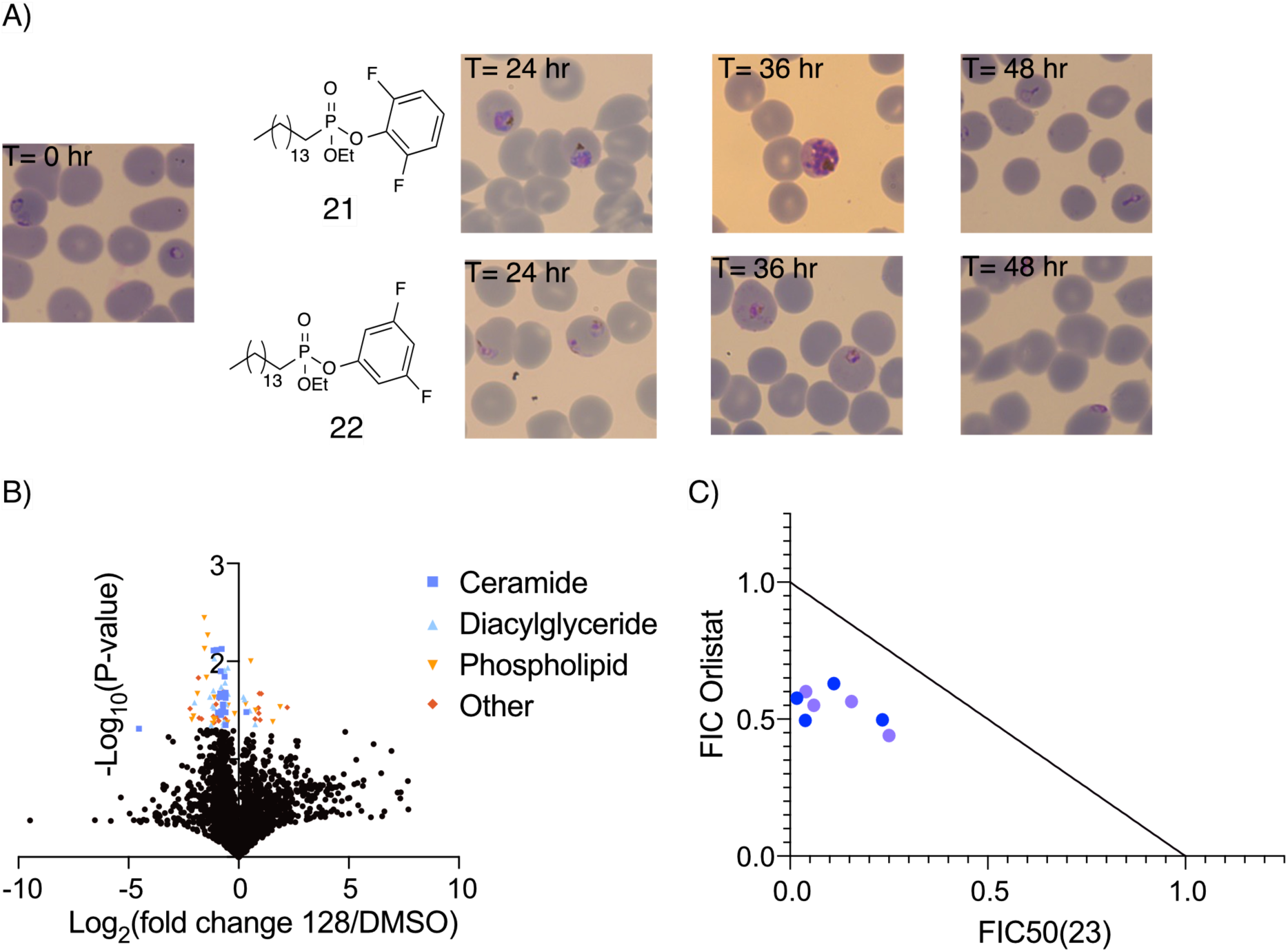
Compound 22 has a mechanism of action different from the pan lipase inhibitor Orlistat. (A) Giemsa-stained parasites. Synchronized ring-stage parasites were treated at 0-6 hours post-invasion with DMSO (vehicle control), 10 µM compound 21 or 22, and imaged at 24 h, 36 h, and 48 h. (B) Lipidomic analysis of compound 22 treated parasites. Synchronized ring stage parasites were treated with either 900 nM compound 22 or DMSO for 24 hours before lysis and collection of lipids. Significantly changed lipid classes are denoted on volcano plot. N= 3-4 (C) Isobolograms depicting P. falciparum in vitro culture-based fractional EC_50_ (FIC50) values in the presence of compound **22** and Orlistat, each point represents a different biological replicate done in technical duplicate.

We next performed metabolomic studies of compound **22** treated parasites to determine if specific metabolic changes occur upon compound treatment. In contrast to Orlistat, which induces accumulation of triglycerides and arrest of parasites in the late schizont stage (Gulati et al., 2015), we did not observe any significant accumulation of individual lipid classes in cells treated with **22** compared to vehicle-treated cells. However, there was an apparent downregulation of both ceramides and diacylglycerides in cells treated with **22** (Figure 5B). However, our efforts to rescue parasites with ceramides and cell-permeable diacylglyceride supplementation were unsuccessful (Figure S6D), suggesting that **22** may cause early mis-regulation of lipid metabolism pathways essential for the synthesis of neutral lipids such as ceramides and diacylglycerides. Nevertheless, our results suggest that **22** targets a pathway distinct from Orlistat. To further confirm this, we conducted drug synergy assays by co-treatment with **22** and Orlistat. These results confirmed a synergistic effect (FIC_50_<1), suggesting that Orlistat and **22** likely target distinct metabolic pathways and inhibition of these pathways can synergize to increase parasite killing (Figure 5C).

### Resistance mediated by compound 22 is mild and similar in mechanism to SalA resistance

Given the similarities in structures between SalA and **22**, we attempted to study potential cross-resistance between SalA and **22** and assessed the relative susceptibility of this compound to induce resistance. Cross-resistance screening of SalA selected parasite lines that we previously generated in Dd2-Polδ parasites (Yoo et al., 2020; Kümpornsin *et al*., 2023) revealed that both P102L and 145I mutations in the PRELI domain-containing protein conferred a similar reduction in susceptibility to **22** compared to that reported for SalA (Figure 6A; 6C). Compared to the Dd2-Polδ parental line, PRELI mutant parasites also showed a 7-10-fold increase in EC_50_ for **22**, suggesting that there may be a common genetic mechanism of resistance.

**Figure 6:**
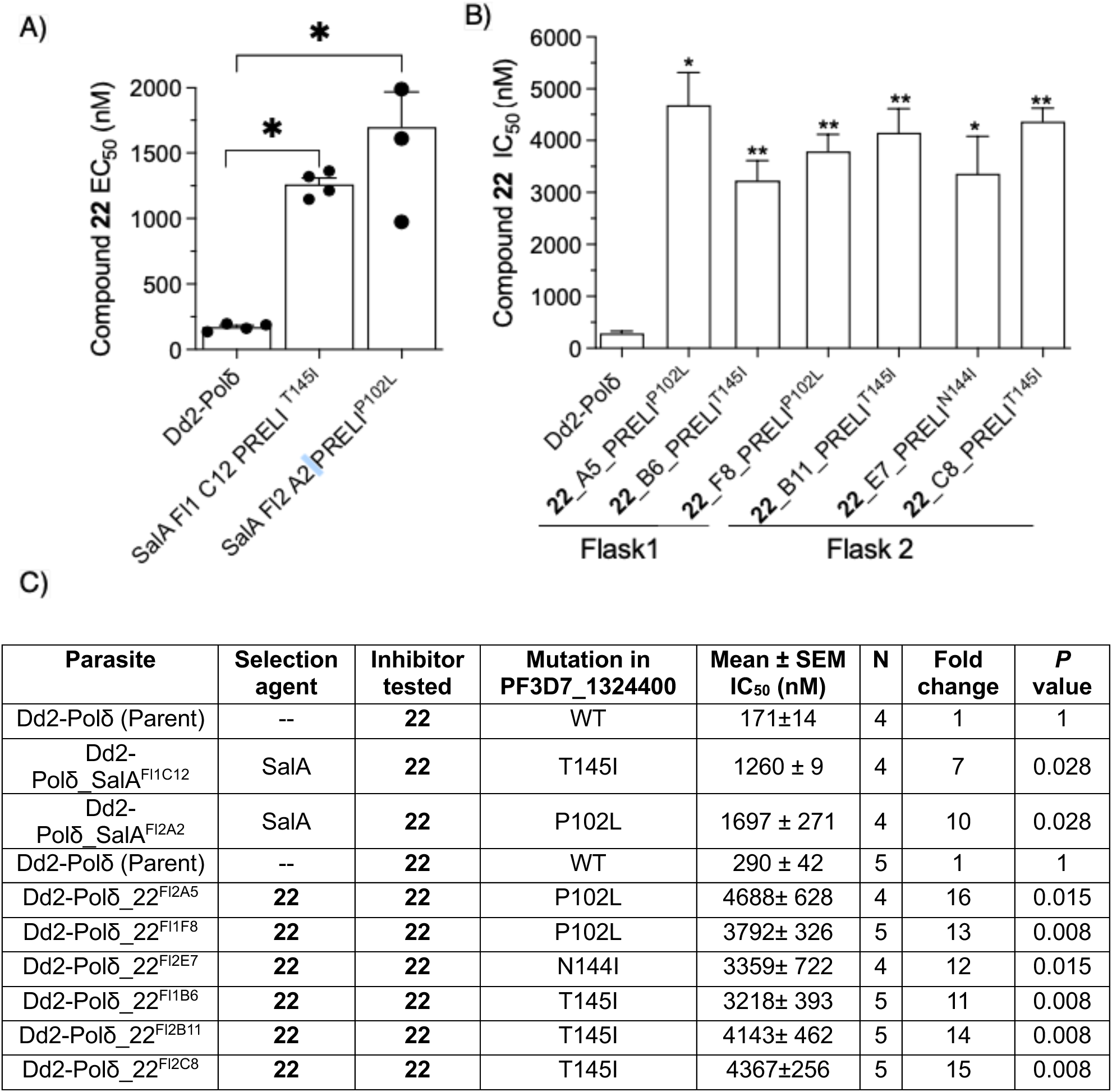
Cross resistance and *in vitro* selection of parasites. (A) Plot of EC_50_ values for two SalA resistant lines (harboring mutations in PRELI domain-containing protein at positions P102L and N145I) for compound **22**. Values are plotted as means ± SEMs and statistical significance determined using Mann-Whitney *U* tests where \**p* < 0.05. (B) EC_50_ shifts of mutant and parental lines against **22**. EC_50_s for each line are presented as means ± SEMs (N, n = 4, 2) and statistical significance determined using Mann-Whitney *U* tests comparing the parental data vs the mutant line where \**p* < 0.05. (C) Summary table of EC_50_s for each line are presented as means ± SEMs (N, n = 4, 2). EC_50_s for each line are presented as means ± SEMs (N, n = 4, 2) and statistical significance determined using Mann-Whitney *U* tests comparing the parental data vs the mutant line where \**p* < 0.05.

To further study the mechanism of resistance to **22** we performed *in vitro* single step selection on the hypermutator Dd2-Polδ parasite line. We pressured 10^9^ asexual blood stage parasites per flask, with duplicate flasks, at a continuous concentration of 5xEC_50_ of compound **22**. We observed recrudescence microscopically in both flasks at day 17 and were able to recover specific clones by limiting dilution. Phenotypic analysis of cloned parasites in 72 hour dose-response assays showed an 11 to15-fold shift in the EC_50_ values of **22** compared with the parental line (Figure 6B).

To identify the mechanism of resistance, we performed whole genome sequencing on eight resistant clones and their parental Dd2-Polδ line. Thia analysis yielded good quality reads corresponding to a high depth of coverage for the resistant clones (Supplemental Table 2). No copy number variations (CNVs) were found in any of the recrudescent parasite genomes relative to 3D7-A in either sequencing experiment except for FL2_A5 which had CNV gain of 1.5 fold in the amplified region of 82kb containing MDR1 (Supplemental Table 2). Analysis revealed that compared to the parental line, resistant clones from both flasks possessed either a P102L, N144I or a T145I mutation in the PRELI domain-containing protein. The P102L and T145I mutations are the same mutations observed in SalA selections (Yoo *et al*., 2020), with N144I being new to selections with **22**.

To determine the minimum inoculum of resistance (MIR), which is an indirect measure of the propensity to generate resistance in the field, we exposed Dd2-B2 parasites to **22** at 5xEC_50_. The selection had a starting inoculum of 3.3 × 10^8^ parasites per flask in three separate flasks (covering the range from 3.3 × 10^8^ to a total of 1 × 10^9^). Parasites recrudesced on day 23 in one out of three flasks, resulting in an MIR value of 1x 10^9^ (Supplementary Table 3). Based on prior observations, drugs with a MIR ≥ 10^9^ have a much lower chances of treatment failure (Duffey et al., 2021). Based on these results, compound **22** is less prone to induce resistance compared to other commonly used antimalarials.

## Discussion

Previously, we used the natural product SalA to identify several α/β serine hydrolases that, when inhibited, caused parasite death in *P. falciparum*. In this study, we describe the synthesis of mixed alkyl/aryl phosphonates as bioisosteres of SalA and further characterize their inhibition of serine hydrolases and antiparasitic effects. We identified a pair of constitutional isomers that enabled a comparative proteomic approach in which targets responsible for the parasite killing effects of the active analog can be identified. Using these compounds as probes, we identified abH112, a previously annotated essential enzyme, that when inhibited in combination with other serine hydrolases, leads to strong anti-parasitic activity of the active compound. Our compound exhibited different stage-specific killing compared to SalA and the pan-lipase inhibitor Orlistat, and the metabolic changes under treatment with our compound differed significantly from previously explored mechanisms of action. This unique phenotype is likely due to the inhibition of abH112, which was not identified as a target of SalA. By continuing to improve our understanding of how serine hydrolases regulate lipid metabolic pathways in *Plasmodium* parasites it is possible to further validate novel drug targets and mechanisms of action that disrupt these pathways.

Previous studies have attempted to assess the importance of essential serine hydrolases using either complete or catalytic inactivation of the enzymes of interest. However, in these cases, either the knockout could not be achieved, or there was insufficient evidence to determine whether the catalytic activity of the enzyme was eliminated (Davison et al., 2022; Yoo et al., 2020). These findings all point to an essential role for metabolic serine hydrolases in the parasite lifecycle. To further validate the essentiality of abh112, we further analyzed a recently reported parasite line in which tetracycline was used to induce a genetic rearrangement in which only the catalytic Ser is mutated to Ala, this keeping expression of the protein but inactivating the enzyme (Davison et al., 2022). We found that the loxP diCre system does in fact enable integration and proper excision of the wild type gene, but we also found evidence that the wild type gene was present in all samples. This suggests that the gene, after being excised, was likely reintegrated into the genome leading to expression of the wild-type protein. This would account for the lack of phenotypic changes associated with the conditional inactivation mutants. Moreover, this provides further support beyond the transposon mutation screens, that abH112 is essential.

Given that conditional knock down of abH112 did not result in parasite growth inhibition or change in potency of compound **22**, it is likely that our inhibitor acts via a pleiotropic mechanism, similar to the parent compound, SalA, on which its design was based. Interestingly, both compound **21** and **22** targeted the essential hydrolase PfMAGL with both acting as pM inhibitors of the recombinant enzyme. However only compound **22** kills parasites suggesting that the combination of inhibition of abH112 and other hydrolases, such as pfMAGL, that results in parasite death. For SalA, which also potently inhibits PfMAGL, parasite killing is likely mediated by inhibition of PfMAGL plus other essential serine hydrolase targets such as the BEM46-like protein (PBLP) or the exported lipase 2 (PfXL2). This leads to a different mechanism of action for SalA and compound **22**. Specifically, we determined that **22** kills parasites at a much earlier stage than previously studied serine hydrolase inhibitors, likely attributed blockage of distinct points in essential metabolic pathways (Gulati et al., 2015; Yoo et al., 2020). Furthermore, we found that **22** synergizes with the pan-lipase inhibitor Orlistat, suggesting that additional serine hydrolase and lipid metabolic pathways could be co-inhibited to increase compound potency and perhaps reduce chances for resistance even further. These avenues for the development of serine hydrolase inhibitors are especially promising, considering that mammalian serine hydrolases can typically be targeted with minimal cytotoxic effects (Bachovchin and Cravatt, 2012). Given the substantial work on the development of specific serine hydrolase inhibitors for human targets with favorable pharmacokinetics and bioavailability, combinations of serine hydrolase inhibitors initially developed for human serine hydrolases could potentially be repurposed and serve as a valuable reservoir for the development of antimalarials.

We also sought to characterize the serine hydrolases targeted by compound **22** using resistance selections, which yielded parasites with only moderate resistance to our compound. We isolated six clones with decreases of potency for compound **22** ranging from 11x-15x compared to the parent parasite line and performed whole-genome sequencing on these clones. Only the PRELI domain-containing protein was mutated in every clone sequenced. The PRELI domain-containing protein is not a serine hydrolase but was also found to be mutated in all clones of SalA-resistant parasites (Yoo et al., 2020). This protein is found in a wide variety of eukaryotic organisms and localizes to mitochondria where it regulates lipid accumulation by shuttling phospholipids across the intermembrane space (Miliara et al., 2019). Previously, the PRELI domain-containing protein was characterized as regulating a mechanism of multidrug resistance in the related *Toxoplasma gondii* parasite (Jeffers et al., 2017). Our results indicate that it likely functions similarly in *P. falciparum* to mediate a mechanism of multidrug resistance, as we demonstrated that compound **22** and SalA likely have different mechanisms of actions and therefore do not directly act on the PRELI domain-containing protein as a primary target. The MIR value for **22** is high and therefore it has high barrier to resistance. Mechanism of action is not always tied to the mechanism of resistance. For example, ART is well established as a drug with a pleiotropic mechanism of action, but resistance to ART is governed by mutations in the k13 gene (Straimer et al., 2015). Moreover, in *Plasmodium* and other organisms, ABC and other efflux pumps are often mutated or upregulated to enable cellular export of drugs with various mechanisms of action (Kim et al., 2021). Ultimately, we did not observe any mutations in serine hydrolases in our resistance selections, which suggests the parasite has difficulty in evolving resistance through mutations in the direct targets of the compounds. Furthermore, the fact that all the identified serine hydrolase inhibitors with potent anti-parasite activity seem to function by inhibition of multiple essential enzymes, may explain why these compounds all have a low propensity to generate resistance in the parasite. These findings further support targeting multiple serine hydrolases to develop new classes of antimalarial drugs or using existing serine hydrolase inhibitors with non-overlapping targets in the parasite to generate therapeutics that prevent resistance. This strategy is also not likely limited to *Plasmodium* parasites, as recent studies in *Staphyloccus auerus* showed that the pleiotropic targeting of serine hydrolases along with other metabolic enzymes leads to potent killing of the bacteria (Bakker et al., 2023).

Here, we have described the use of mixed alkyl/aryl phosphonates as a nascent class of serine hydrolase inhibitors with potent antimalarial properties. These compounds have underscored the value of target-agnostic screening to identify new drug susceptible pathway and have the potential for application in other diseases and pathogens where serine hydrolases also play essential roles in maintaining homeostasis (Babin et al., 2022; Li et al., 2022). Mixed alkyl/aryl phosphonates are synthetically tractable and provide a starting point for the development of molecules that enable the validation of hydrolases that are optimal drug targets that can be targeted simultaneously. The continued development of chemical tools to study lipid metabolizing pathways and proteins governing such pathways will be essential for the development of next-generation therapeutics for *P. falciparum*. Furthermore, future studies will likely benefit from combining phenotypic screening, target identification, and characterizations of changes in metabolism to prioritize the critically important serine hydrolases as candidate drug targets.

### Significance

Lipid metabolism and the dynamic regulation of lipid membranes are essential pathways within *P. falciparum*. Previously we used the potent antimalarial natural product Salinipostin A to identify a series of serine hydrolases which, when inhibited, results in parasite death. Here, we developed mixed alkyl/aryl phosphonates as tool compounds to aid in dissecting and identifying important regulators of lipid metabolism. We described the use of a pair of constitutional isomers to identify a serine hydrolase, abH112, as a possible new drug target. Moreover, we showed how a previous attempt of a conditional knockout of abH112 led to gene duplication and that further methodologies are needed to investigate essential lipid metabolizing enzymes within the parasite. We also described a mechanism of killing not previously identified with SalA or the pan-lipase inhibitor Orlistat, opening up the possibility of multiple lipid metabolizing pathways that can be targeted in drug development. Lastly, we demonstrated that PRELI domain-containing protein is likely a mechanism of multidrug resistance with mutations found in it rather than targeted serine hydrolases, similar to SalA. Ultimately, we have described how mixed alkyl/aryl phosphonates can be used to study lipid metabolizing pathways and further bolstered serine hydrolases as an important class for antimalarial drug development.

## Acknowledgments

This work was supported in part by the NIH (R33 AI127581, to DAF and MB, R01 AI109023 to D.A.F) and by Department of Defense grant (W81XWH2210520 to D.A.F., M.B). VLL was supported by NIH training grant T32 GM113854. The work was also supported by funding to MJB from the Wellcome Trust (220318/A/20/Z) and the Francis Crick Institute (https://www.crick.ac.uk/) which receives its core funding from Cancer Research UK (CC2129), the UK Medical Research Council (CC2129), and the Wellcome Trust (CC2129).

## Author Contributions

Conceptualization, J.M.B and M.B; Methodology, J.M.B, S.K.N., S.K., D.F., M.B.; Formal Analysis, J.M.B, S.K.N., S.K, T.Y.; Investigation, J.M.B, S.K.N, S.K., D.A., F.H., T.Y., V.L.. R.M., F.F., S.L.; Writing-Original Draft, J.M.B., S.K.N., D.F., M.B.; Writing-Editing and Revising, J.M.B., S.K.N., D.F., M.B; Visualization, J.M.B, S.K.N., Supervision, M.B., A.A., E.Y., D.F., M.B.; Funding Acquisition, D.F., M.B.

## STAR Methods

### *P. falciparum* Parasite Culture

*P. falciparum* W2, 3D7, and Dd2_Polδ cultures were maintained, synchronized, and lysed as previously described (Straimer et al., 2017). Parasite cultures were grown in human erythrocytes (no blood type discrimination) purchased from the Stanford Blood Center (approved by Stanford University) or the Interstate Blood Bank (for Columbia University). at 3% hematocrit in RPMI-1640 media, supplemented with 25 mM HEPES, 50 mg L-hypoxanthine, 2mM L-glutamine, 0.225% sodium bicarbonate, 0.5% (wt/vol) AlbuMAXII (Invitrogen) and 10 μg/mL gentamycin. Cultures were maintained at 37°C in modular incubator chambers (Billups-Rothenberg) at 5% O2, 5% CO2 and 90% N2.

### PfMAGL and hMAGL Inhibition Assays

5 µL of 20 nM PfMAGL or hMAGL (final concentration 5 nM) in PBS containing 0.1% Triton was preincubated with 5 µL of inhibitor diluted in PBS for 30 minutes at 37°C in a 384-flat black well plate. 10 µL of 80 µM 4-methylumbelliferyl caprylate (final concentration 40 µM) in 1% DMSO in PBS was then added and fluorescence (lex = 365 nm and lem = 455 nm) was read at 37°C in 1 min intervals on a Cytation 3 imaging reader (BioTek, Winooski, VT, USA) for 60 min. Turnover rates were calculated over the linear phase of the reaction in RFU/min.

### *P. falciparum* Replication Assay

Synchronized ring stage parasites culture at 1% parasitemia and 0.5% hematocrit were plated in a 96-well plate and dilutions of compounds in 1xPBS were added to parasites. The culture was incubated for 72 hours before the samples were fixed in a final concentration of 1% paraformaldehyde in PBS for 30 minutes at room temperature. The fixed samples were then stained with a 50 nM final concetration of the YOYO-1 nuclear stain. Parasite replication was monitored by observing either YOYO-1 positive red blood cells (infected) or YOYO-1 negative cell (unifected) using a BD Accuri C6 automated flow cytometer. For one hour pulse assay, cultures were treated with drug for one hour before being washed and resuspended in fresh media, and then grown for an additional 71 hours. Rescues were performed with concentration of lipid added at beginning of assay.

### *P. falciparum* Rate of Kill Assay

Parasite killing kinetics was adapted from the method previously outlined in previously (Linares et al., 2015). In brief, 3D7 ring-stage parasites at 0.5% parasitemia and 2% hematocrit and were treated with 10 x EC_50_ of drug. At either 24 hours or 48 hours, drug free parasites were then cultured in fresh erythrocytes and culture media for the remaining time until readout at 72 hours post treatment. Cultures were then fixed and stained as above.

### Human Cell Cytotoxicity Assay

Human Foreskin Fibroblasts in Dulbecco’s modified Eagle’s medium (DMEM) with 10% FetalPlex animal serum complex (Gemini Biotech, catalog no. 100602) were seeded at 2500 cells (nonconfluent) per well <24 h before addition of the compound. Compounds were diluted for dose−response concentrations in media and added to the cells for 72 h. Cell viability was measured using the CellTiter-Blue Assay (Promega) as per the manufacturer’s instructions.

### Preparation of P. falciparum Lysates for Proteomics

75 mL cultures of W2 parasites at 12-15% parasitemia in 4% hemotocrit were either pretreated with 0.2% DMSO in PBS, 5µM Compound 21, or 5µM Compound 22 for 2 hours at 37°C. Then either DMSO, 5µM 21-alk, or 22-alk was added to the cultures and incubated for another 2 hours at 37°C. Labelled parasites pellets were harvested via saponin lysis and pellets were lysed in 0.1% SDS in PBS using probe sonication on ice.

### Competitive LC-MS/MS target identification

Protein concentration was measured using the Bradford assay and 250 µg of samples (0.5 mg/mL, 500 µL) were used. Click chemistry was performed for one hour at r.t. with the addition of biotin azide (50 µM, 50x stock in DMSO), tris(2-carboxyethyl)phosphine hydrochloride (TCEP) (1 mM, 50x fresh stock in water), tris[(1-benzyl-1H-1,2,3-triazol-4-yl)methyl]amine (TBTA) (100 µM, 16x stock in DMSO:tButanol 1:4), and copper(II) sulfate (1 mM, 50x stock in water). Protein was precipitated by adding 4 vol. −20 °C acetone and samples were incubated overnight at −20 °C. Samples were centrifuged for 10 min at 20’000 x g at 4 °C yielding a protein pellet which was air dried for 20 min. Pellets were solubilized in 2% SDS in PBS via sonication. Tubes were centrifuged at 4’700 x g for five min and the soluble fraction was transferred to a new tube. PBS was added to give a final SDS concentration of 0.2%. 60 µL of streptavidin agarose beads (ProteoChem) were added and the mixture was rotated for four hours at r.t. Beads were washed with 1% SDS in PBS (1x 10 mL), PBS (3× 10 mL), and water (3x 10 mL). Beads were resuspended in 6 M urea in PBS (500 μL), reduced with 10 mM neutralized TCEP (20x fresh stock in water) for 30 min. at r.t., and alkylated with 25 mM iodoacetamide (400 mM fresh stock in water) for 30 min. at r.t. in the dark. Beads were pelleted by centrifugation (1’400 x g, two min.) and resuspended in 150 μL of 2 M Urea, 1 mM CaCl_2_ (100x stock in water), and trypsin (Thermo Scientific, 0.5 μL of 0.5 μg/μL) in 50 mM NH_4_HCO_3_. The digestion was performed overnight at 37 °C. Samples were acidified with 1 vol. of isopropanol 1% TFA and desalted using styrenedivinylbenzene reverse-phase sulfonate (SDB-RPS) StageTips as described previously (Brunner et al., 2022). Briefly, samples were loaded on a 200 µL StageTip containing two SDB-RPS disks and centrifuged at 1500 x g for 8 min. This was repeated until all the sample was loaded on the StageTip. StageTips were washed three times with 200 µL of isopropanol 1% TFA at 1500 x g for 8 min, then eluted with 100 µL of 80% MeCN, 19% water, and 1% ammonia and dried. Samples were then analyzed by LC-MS/MS, MaxQuant, and Perseus.

### LC-MS/MS analysis

Peptides were resuspended in water with 0.1 % FA and analyzed using an EASY-nLC 1200 nano-UHPLC coupled to a Q Exactive HF-X Quadrupole-Orbitrap mass spectrometer (Thermo Scientific). The chromatography column consisted of a 50 cm long, 75 μm i.d. microcapillary capped by a 5 μm tip and packed with ReproSil-Pur 120 C18-AQ 2.4 μm beads (Dr. Maisch GmbH). LC solvents were 0.1 % FA in H_2_O (Buffer A) and 0.1 % FA in 90 % MeCN: 10 % H_2_O (Buffer B). Peptides were eluted into the mass spectrometer at a flow rate of 300 nL/min. over a 90 min linear gradient (5-35 % Buffer B) at 65 °C. Data was acquired in data-dependent mode (top-20, NCE 28, R = 15,000) after full MS scan (R = 60,000, m/z 400 – 1,300). Dynamic exclusion was set to 10 s, peptide match to prefer and isotope exclusion was enabled.

### MaxQuant analysis

The MS data were analyzed with MaxQuant and searched against the Plasmodium falciparum proteome (isolate 3D7, Uniprot) or human proteome (Uniprot) and a common list of contaminants (included in MaxQuant)(Cox et al., 2014). The first peptide search tolerance was set at 20 ppm, 10 ppm was used for the main peptide search and fragment mass tolerance was set to 0.02 Da. The false discovery rate for peptides, proteins, and site identification was set to 1 %. The minimum peptide length was set to 6 amino acids and peptide re-quantification and “match between runs” were enabled. MaxLFQ without normalization was used. Methionine oxidation and N-terminal acetylation were searched as variable modifications and carbamidomethylation of cysteines as fixed modification.

### Phylogenetic and Structural Model

The human, Taxoplasma gondii, and Plasmodium falciparum proteomes were filtered for proteins annotated to be in the Hydrolase 4 (PF12146) protein family domain. All sequences for these proteins were input into the Clustal Omega algorithm for multiple sequence alignments and a dendrogram was constructed based on homology. For structural models, the AlphaFold structures of both ABHD17a and abH112 were placed into Pymol and the alignment feature was used.

### Stage Specific Killing Assay

Tightly synchronized W2 ring stage cultures at 5% parasitemia in 2% hematocrit were treated with either 10 µM compound 21 or 22. Every 12-24 hours, a blood smear was taken from the cultures and visualized with 5% giemsa stain to monitor the growth of the parasite.

### Preparation of Lipidomic Samples

For lipid analysis, synchronized W2 ring stage of 50 mL cutlures of 10% parasitemia in 4 % hematocrit were treated with 920nM (5x IC_50_) of compound 22 or DMSO vehicle for 16 hours. Red blood cells were saponinin lysed and the parasite pellet was collected. Parasite pellets were subsequently collected and resuspended in 1 ml of 1xPBS buffer and transferred into glass vials pre-loaded with 2:1 chloroform/methanol to a final ratio of 2:1:1 chloroform/methanol/1xPBS. Samples were vigorously shaken spun down at 1000g and the organic layer was collected. Three vials of lipids extracted from each treatment were dried down under a stream of argon.

### Lipid analysis using high-performance liquid chromatography-mass spectroscopy

Lipidomics was performed on an Agilent 6545 Q-TOF LC/MS as previously described and data was anaylzed using the online XCMS platform (Onguka et al., 2021). Metabolites shown to be significantly changed between the two treatment groups were then searched by their mass and classified using the online LipidMaps tool (Conroy et al., 2023).

### Food Vacuole Visualization

Parasites were plated at 5% parasitemia, 4% hematocrit in a 6 well plate and treated with either DMSO, **21,** or **22** at 5 µM for 16 hrs. Nile Red was then added to a final concentration of 1 μg/ml into the parasite culture and the cultures were incubated on ice for 30 min. The culture subsequently washed with 1xPBS before analysis by confocal microscopy (excitation/ emissions maxima is 552/636 nm). The resulting images were then analyzed using ImageJ and the area of food vacuole was measured by circling the Nile Red positive areas.

### Drug Synergy Assay

Synchronized ring stage W2 parasites culture at 1% parasitemia and 0.5% hematocrit were plated in a 96-well plate. Drug mixtures in four different ratios diluted in PBS were added to the cultures and the parasites were grown for 72 hours. EC_50_ values were derived for each compound tested alone, and fractional EC_50_ (FIC50) values were determined for each compound tested in combination (FIC50 =EC_50_ of the drug alone/EC_50_ of the drug in combination) and plotted for each drug combination.

### *In vitro* drug selection studies

*P. falciparum* hypermutator (Dd2-Polδ) was used for *in vitro* drug selection. To select for compound **22** resistant parasites, duplicate flasks of 2×10^9^ parasites were exposed to continuous drug pressure at concentrations of 5xEC_50_ for 60 days until recrudescent parasites were observed. Drug media was changed daily for first 6 days. Cultures were monitored daily by microscopic examination of Giemsa-stained smears. Recrudescent cultures were assayed to analyze degree of resistance and the resistant parasites were obtained by limiting dilution. Dd2-Polδ line was maintained in culture throughout the *in vitro* selection.

### Minimum inoculum of resistance studies

To determine MIR, 3.3×10^8^ Dd2-B2 parasites were cultured per flask in triplicates continuous under 5xEC_50_ drug pressure. Media was changed daily for first 6 days and culture was monitored daily by Giemsa-stained smears for complete killing and recrudescence. Cultures were passaged every week by giving fresh blood and were maintained for 60 days. The MIR is defined as the minimum parasite inoculum used to obtain resistance and calculated as follows: total number of asexual blood stage (ABS) parasites cultured ÷ total number of recrudescent cultures.

### *In vitro* drug susceptibility assays

To define the baseline EC_50_ of Dd2-Polδ to compound **22** and shifts in susceptibility in recrudescent lines, parasite cultures at 0.2% parasitemia and 1% hematocrit (HCT) were incubated for 72 hours with a range of ten drug concentrations that were 2-fold serially diluted in duplicates along with drug-free controls in complete medium. Parasite survival was assessed by flow cytometry on IntelliCyt iQue3 (Sartorius) using SYBR Green and MitoTracker Deep Red FM (Life Technologies) as nuclear and mitochondria stains, respectively. Growth inhibition was plotted using nonlinear regression function to calculate EC_50_ values (using Graphpad Prism 7).

### DNA Extraction and Whole-Genome Sequencing

DNA libraries for each sample were prepared using the Illumina Nextera DNA Flex library kit with dual indices. The samples were multiplexed and sequenced on an Illumina MiSeq to obtain 300 bp paired end reads and Nextseq to obtain 150 bp paired end reads at 19-44x depth of coverage across the samples. The sequence data generated were aligned to the *P. falciparum* 3D7 genome (PlasmoDB version 48.0) and variants filtered based on quality scores and read depth to obtain high quality SNPs. The list of variants from the resistant clones were compared against the Dd2-Polδ parent to obtain homozygous SNPs present exclusively in the resistant clones.

### Generation of abH112 cKD parasites for translation regulation

To generate abh112 cKD parasites, CRISR-Cas9 based TetR-DOZI aptamer system was used to regulate its expression (Murithi et al., 2021). The left homology region fused with recodonized 3’ end of ABH112 and right homology region as well as guide sequence were cloned into pSN054 donor plasmid containing 3XHA epitope tag at C-terminus as described previously(Nasamu et al., 2021).The final construct was transfected into Cas9 expressing NF54 parasites and maintained in 500nM aTC and 2.5 mg/mL of Blasticidin S. Edited parasites were confirmed by site specific integration PCR. All the primers, guide sequence and synthetic fragment sequence are given in supplementary table S1.

### Western blot analysis

Synchronized abH112 cKD ring stage parasites were cultured in presence and absence of aTC for 72 hours. Cultures were harvested and RBCs were lysed using 0.05% saponin in 1xPBS at 4°C. Parasites were washed two times with cold1xPBS containing cOmplete proteases inhibitors (4693159001, Sigma) and pellets were resuspended in SDS buffer (8 M Urea, 5% SDS, 50 mM Bis-Tris, 2 mM EDTA, 25 mM HCl, pH 6.5) and Laemmli buffer. Protein samples were run on 10% Mini-PROTEAN® TGX™ Precast Gels (Biorad) and transferred onto nitrocellulose membranes (Biorad). The membrane was blocked in 3% BSA-TBS and probed with either mouse anti-HA (1: 1000) or rabbit anti-Bip followed by HRP conjugated secondary antibodies. Protein blots were developed using ECL Chemiluminescent Substrate and imaged and analyzed by ChemiDoc^TM^ MP Imaging System (Biorad).

### Asexual growth rate analysis

To assess asexual parasite growth rate, synchronous ring-stage NF54 and abH112 cKD parasites were cultured at different aTC concentrations (30 nM, 3 nM and no aTC) over two intra-erythrocytic developmental cycles. Parasite expansion rate was assessed by flow cytometry on IntelliCyt iQue3 (Sartorius) using SYBR Green and MitoTracker Deep Red FM (Life Technologies) as nuclear and mitochondria stains, respectively.

### General Synthetic Procedures

#### Appel Reaction Alcohol to Iodo

Alcohol (1 equivalent) was dissolved in dichloromethane (final concentration of 0.2 M). To the solution, iodine (1.2 equivalents) and triphenylphosphine (2 equivalents) were added and the reaction was protected from light and stirred for 3 hrs at room temperature. The reaction was quenched with sodium thiosulfate and the aqueous phase was extracted with dichloromethane (2×) and the combined organic phase was washed with brine and then dried over sodium sulfate and concentrated under reduced pressure. The crude reaction mixture was then purified via silica flash chromatography and concentrated under reduced pressure to yield iodo products.

#### Arbuzov Reaction

Iodo containing compound (1 equivalent) is dissolved in triethyl phosphite (4 equivalents) and the solution is stirred at 130°C for 16 hours. The product is concentrated via constant airflow over the solution and the crude product was carried forward.

#### Direct Aryloxylation

Diethyl phosphonate (1 equivalent) was dissolved in dichloromethane (final concentration 0.2 M) and to the solution triflate anhydride (1.5 equivalents) and pyridine (2 equivalents) were added sequentially (the reaction will bubble release fumes and turn orange) and stirred for 10 minutes at room temperature. Various phenols (2.5 equivalents) were then added and the reaction and stirred for 30 more minutes at room temperature. The reaction was then concentrated under reduced pressure and the reaction mixture was purified via silica flash chromatography and concentrated under reduced pressure to yield mixed alkyl/aryl phosphonates.

#### Compound 26

Alcohol (1 g, 1 eq), DMAP (367 mg, 1 eq), and triethylamine (2.5 mL, 6 eq) were stirred at room temperature for 10 min in 30 mL of DCM. Tosyl chloride (1.15 g, 2 eq) was added and the reaction was stirred overnight. The reaction was quenched with cold water, extracted with ethyl acetate 3x, washed with 1M HCl, concentrated NaHCO_3_, brine, dried over sodium sulfate. The residual material was then purified via silica flash chromatography (0-20% ethyl acetate in hexanes) (1.548 g, 96% yield).

#### Compound 27

Lithium Aluminum Hydride (450 mg, 6 eq) was dissolved in dry THF (35 mL). Ester (1.54 g, 1 eq) was dissolved in 3 mL of THF and added dropwise to solution and stirred for 4 hours at 55°C. Reaction was quenched with water, filtered through celite, redissolved in ethyl acetate and dried over sodium sulfate. The residual material was then The residual material was then purified via silica flash chromatography (0-40% ethyl acetate in hexanes) (50% yield).

#### Deprotection of Compound 28

Silyl protected compound (450 mg, 1 eq) and potassium hydroxide (112mg, 4eq) were stirred in methanol at 50°C. The reaction was quenched with water and the pH was adjusted to 1 using 1M HCl and extracted with ethyl acetate 3x, and the combined organic layer was washed with brine and dried over sodium sulfate. The residual material was then purified via silica flash chromatography (0-40% ethyl acetate in hexanes) to yield (400 mg, 97 % yield).

