## Supplemental Methods, Figures and Tables for "Mixed Alkyl/Aryl Phosphonates Identify Metabolic Serine Hydrolases as Antimalarial Targets"

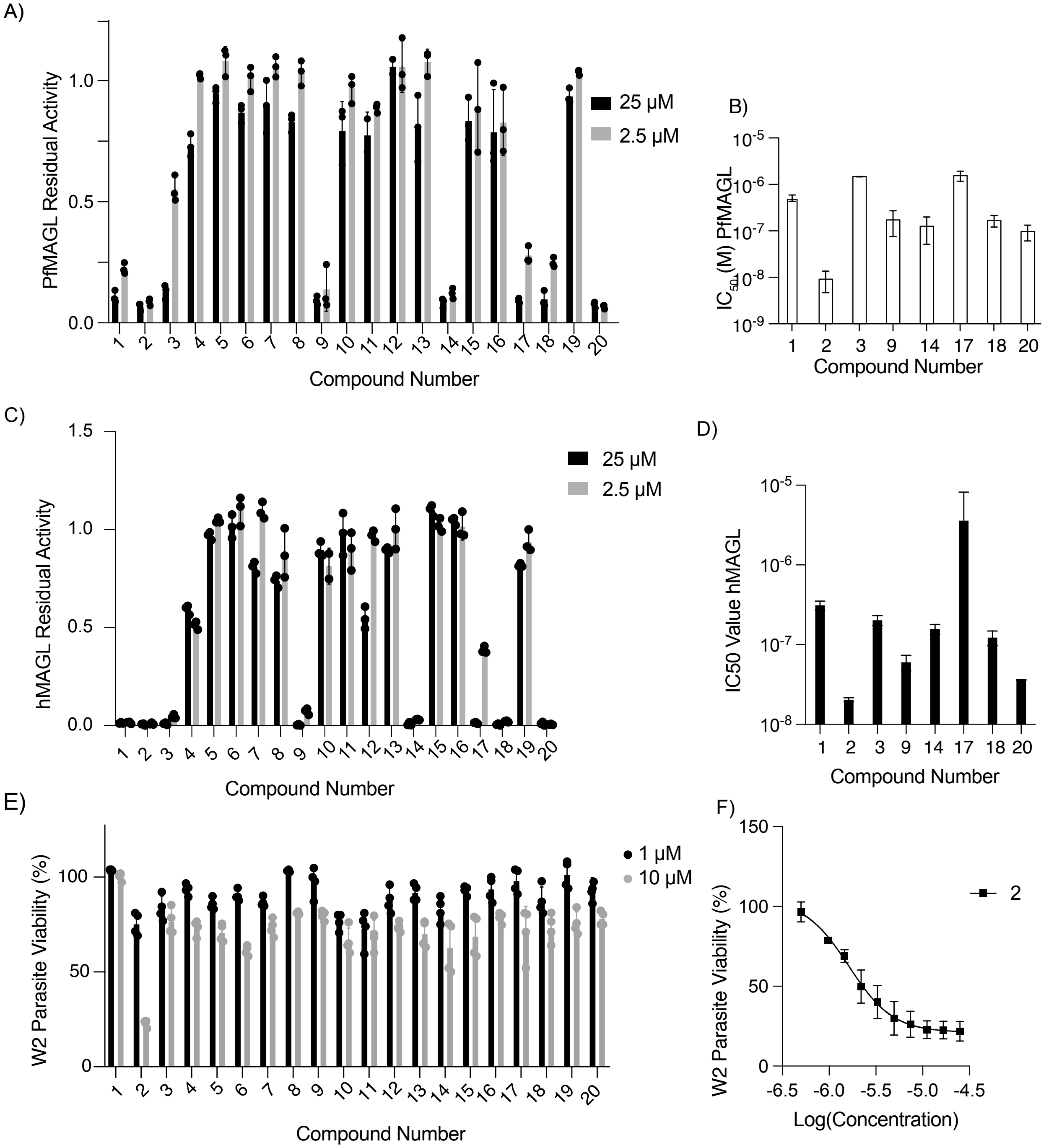


**Figure S1 (Related to Figure 1)**: Biochemical and Phenotypic Characterization of Initial Phosphonate Library.

(A) Screen of phosphonates against PfMAGL at either 25 µM (black) or 2.5 µM (grey) (mean ± SD, N,n = 2,3).

(B) Screen of phosphonates against hMAGL at either 25 µM (black) or 2.5 µM (grey).

(C) $IC_{50}$ values for phosphonates against hMAGL(mean ± SD, N,n = 2,3). D) Dose response curve for W2 parasites treated with varying concentrations of compound 2(mean ± SD, N,n = 3,2).

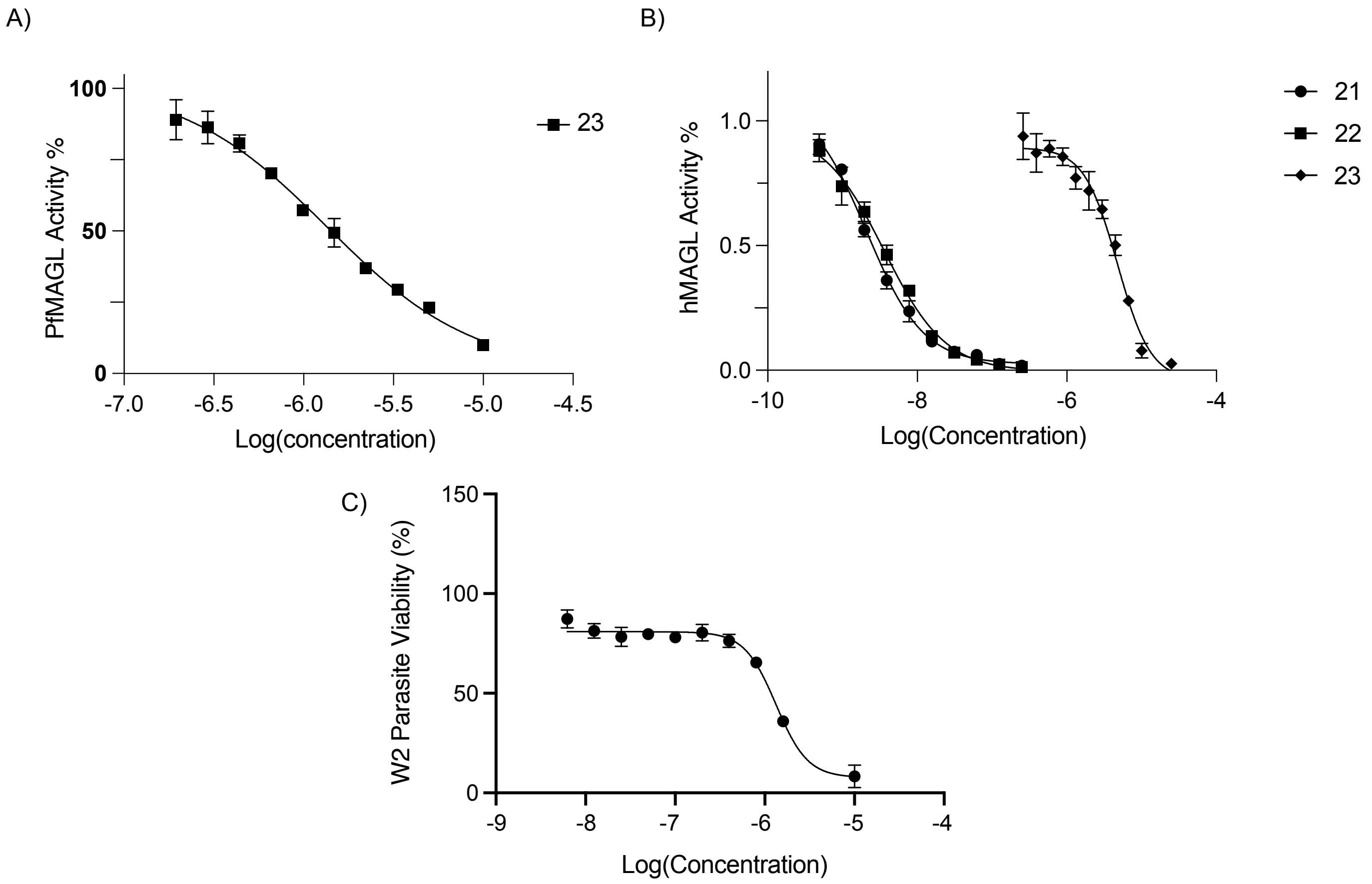


**Figure S2 (Related to Figure 2)**: Characterization of Long Chain Phosphonates.
(A) Dose response curve for PfMAGL treated with varying concentrations of compound **23** (mean ± SD, N,n = 2,3).
(B) Dose response curve for hMAGL treated with varying concentrations of the long chain phosphonates (mean ± SD, N,n = 2,3).
(C) Dose response curve for W2 parasites treated with varying concentrations of compound **2** for one hour before parasites were washed and growth was quantified after 72 hours (mean ± SD, N,n = 4,2).

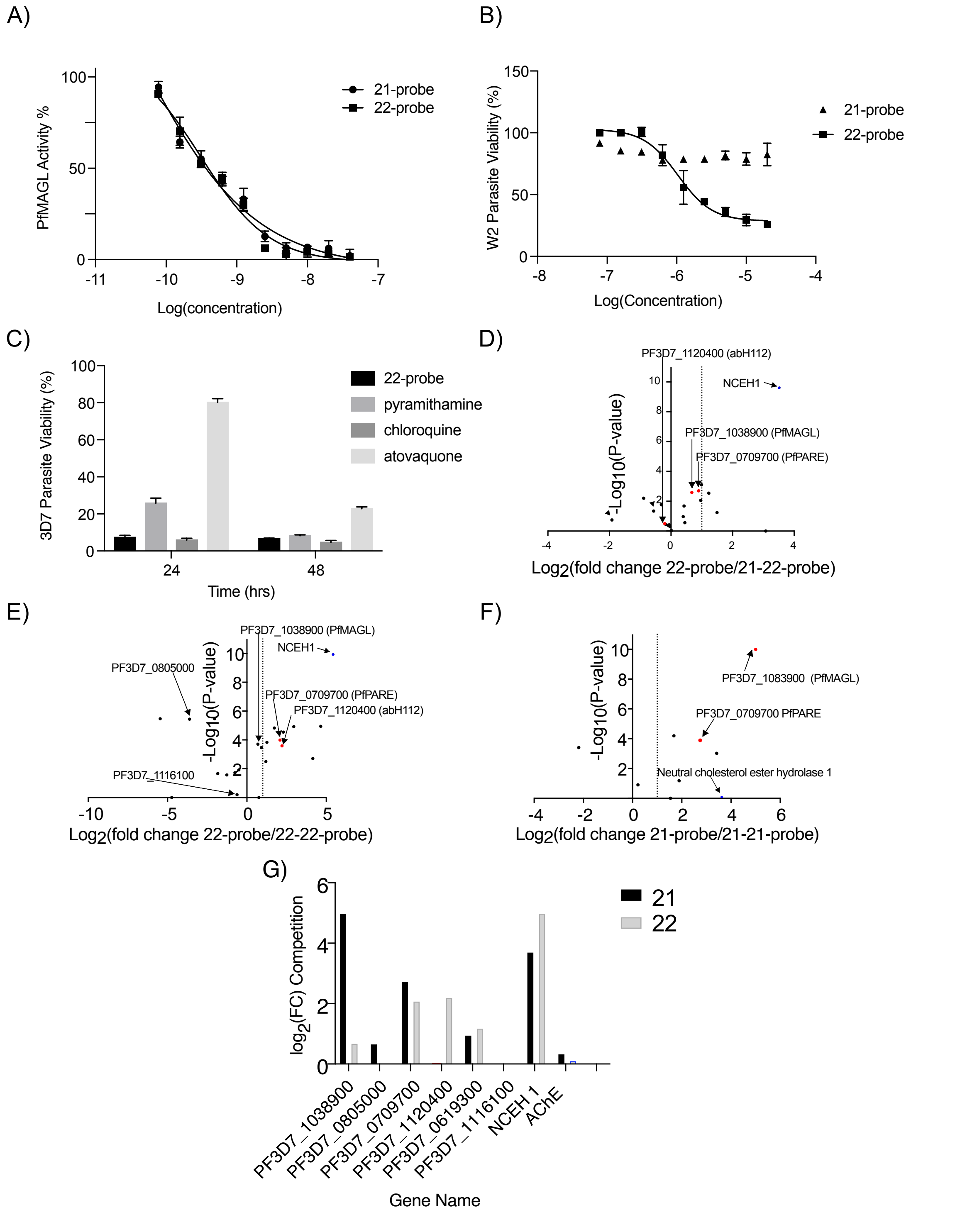


**Figure S3 (Related to Figure 3):** Characterization of Long Chain Phosphonates Probes and Competitive Activity-Based Protein Profiling.
(A) Dose response curve for hMAGL treated with varying concentrations of the long chain phosphonates (mean ± SD, N,n = 2,3).

(B) Dose response curve for W2 parasites treated with varying concentrations of compo for one hour before parasites were washed and growth was quantified after 72 hours (mean ± SD, N,n = 4,2).
(C) Rate of Kill of Compound **22-probe** measured against compared against known fast (chloroquine), medium (pyrimethamine), and slow (atovaquone) acting inhibitors of parasite growth in which compounds were dosed at their $EC_{50}$ and parasite viability was monitored over time and ± SEM. N,*n* = 6,2.
(D) Volcano plots of Activity-based probe **22-probe** targets competed by compound **21** in P. falciparum. The x axis shows the logarithmic base 2-fold change between **22-probe**-treated and **21** pretreated parasites. The statistical significance was determined without correcting for multiple comparisons in triplicates and the y value in the volcano plot is the negative logarithm of the p value. Human serine hydrolases are colored blue and Plasmodium are colored red.
(E) Volcano plots of Activity-based probe **22-probe** competed by compound **22** targets in *P. falciparum*. The x axis shows the logarithmic base 2-fold change between **22-probe**-treated and **22** pretreated parasites. The statistical significance was determined without correcting for multiple comparisons in triplicates and the y value in the volcano plot is the negative logarithm of the p value. Human serine hydrolases are colored blue and Plasmodium are colored red.
(F) Volcano plots of Activity-based probe **21-probe** competed by compound **21** targets in *P. falciparum*. The x axis shows the logarithmic base 2-fold change between **21-probe**-treated and **21** pretreated parasites. The statistical significance was determined without correcting for multiple comparisons in triplicates and the y value in the volcano plot is the negative logarithm of the p value. Human serine hydrolases are colored blue and Plasmodium are colored red.
(G) Quantification of fold change in competitive ABPP experiments.

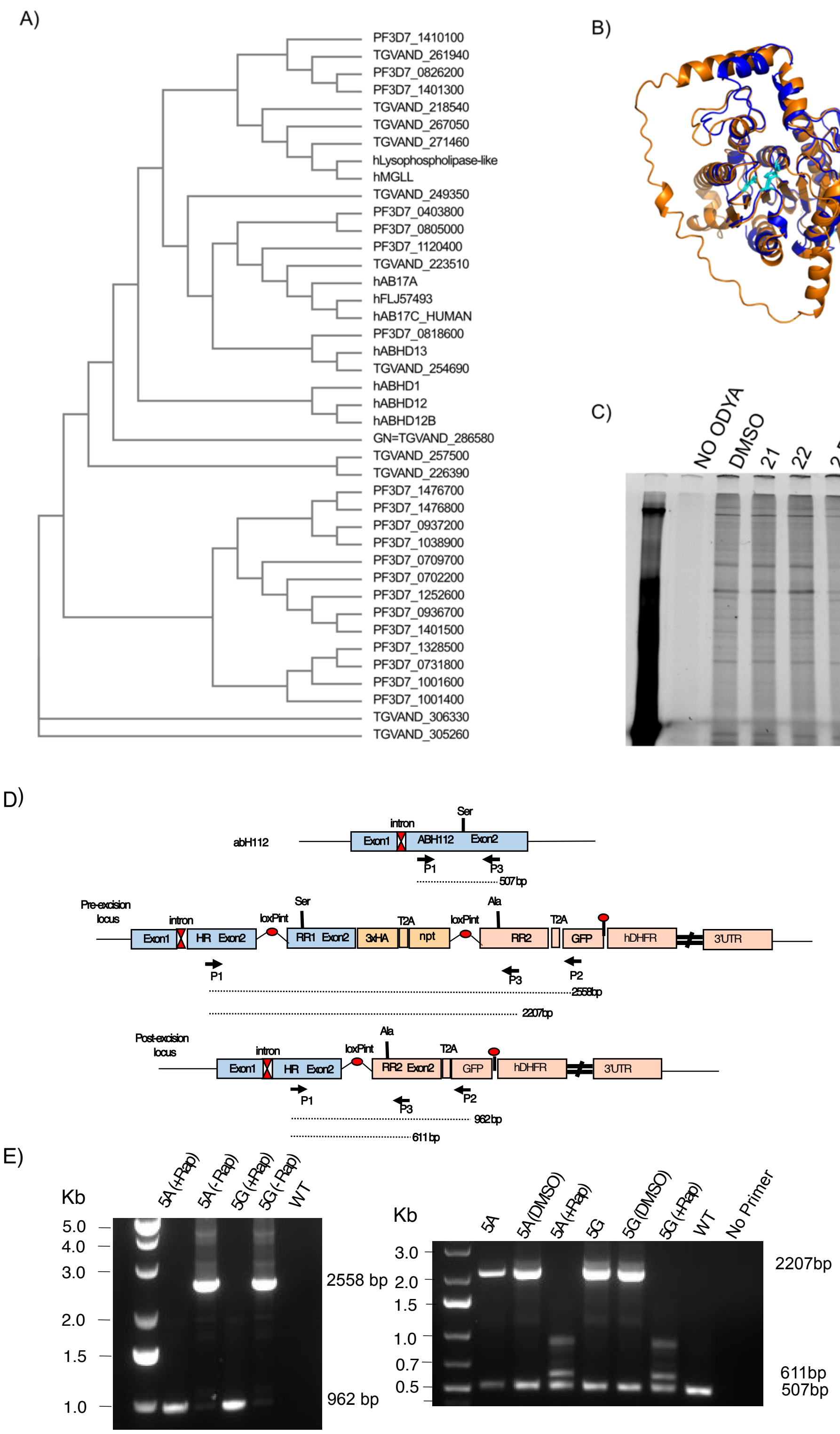


**Figure S4 (Related to Figure 4)**: Catalytic Inactivation of abH112 leads to gene duplication. (A) Dendrogram of *Plasmodium falciparum*, *Toxoplasma gondii*, and human serine hydrolases that share a protein family domain generated using the Clustal Omega algorithm.

(B) Overlay of Predicted AlphaFold structure of abH112 (orange) and hAB17A (blue) with active site catalytic triad in cyan.
(C) Parasites were incubated with 17-ODYA for 16 hours before treatment with either DMSO, Compound 21, 22, or 2-BP and palmitoylation before cell lysates were reacted with TAMRA-azide and visualized using SDS-PAGE.
(D) Schematic of LoxP-diCre system with Wild-type, pre-excision, and post-excision genes
(E) PCR of pre (-Rap) and post excision (+Rap) LoxP parasites with primers P1 and P2 (left) and primers P1 and P3 (right).

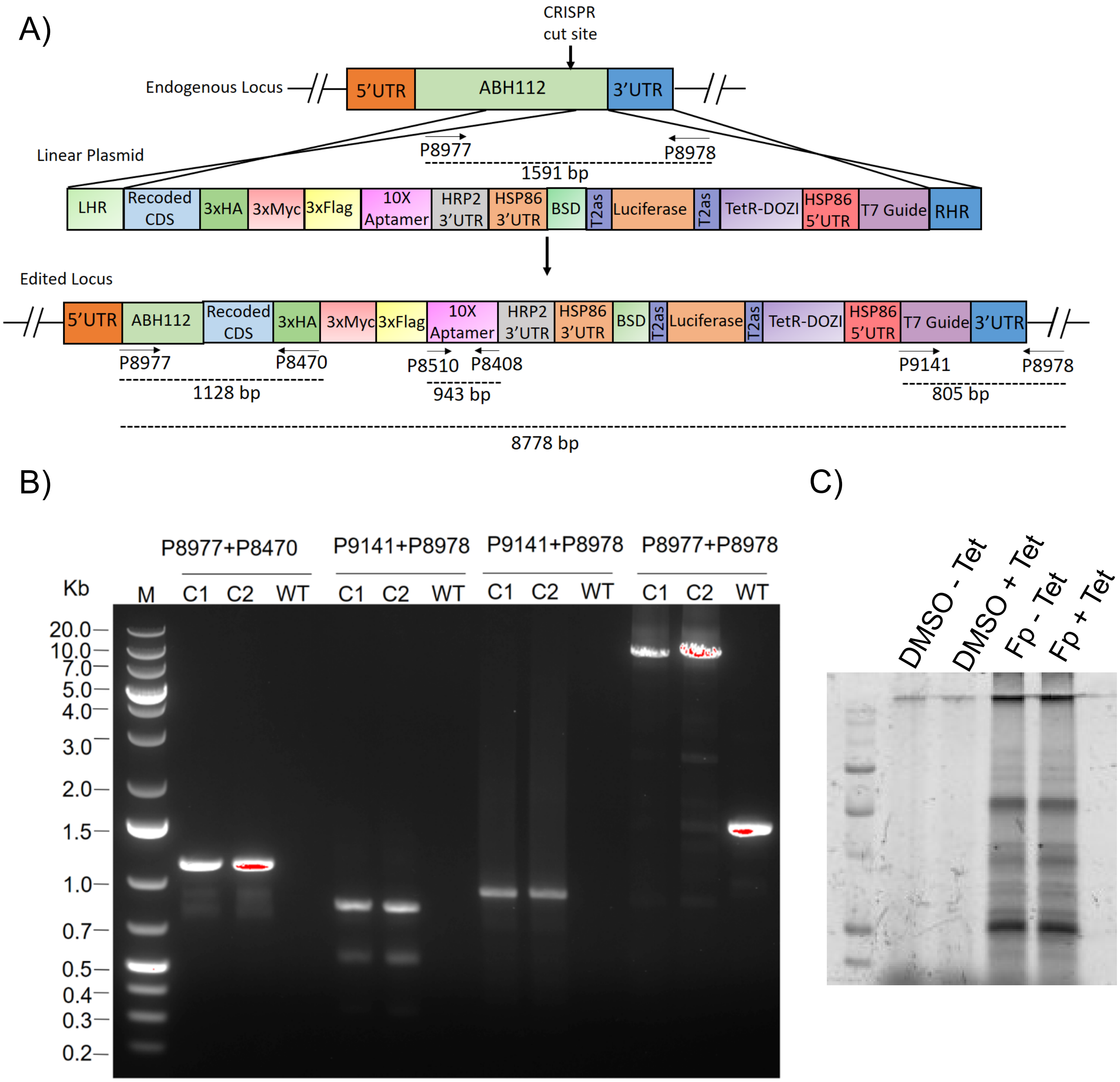


**Fig S5 (Related to Figure 4):** Generation of abH112 cKD lines.

(A) A schematic of the generation of abH112 cKD lines. Primers used for diagnostic PCRs are indicated.

(B) Correct integration was confirmed using primer sets P8977/P8470 and P9141/P8978 designed beyond sites of recombination that amplified 1128 and 805 bp products for 5' and 3' integration respectively. Aptamers were cirmed by primer set P8510/P8408.

(C) Global serine hydrolase activity in conditional knockdown parasites, using Fp-alkyne were reacted with TAMRA-azide and visualized using SDS-PAGE.

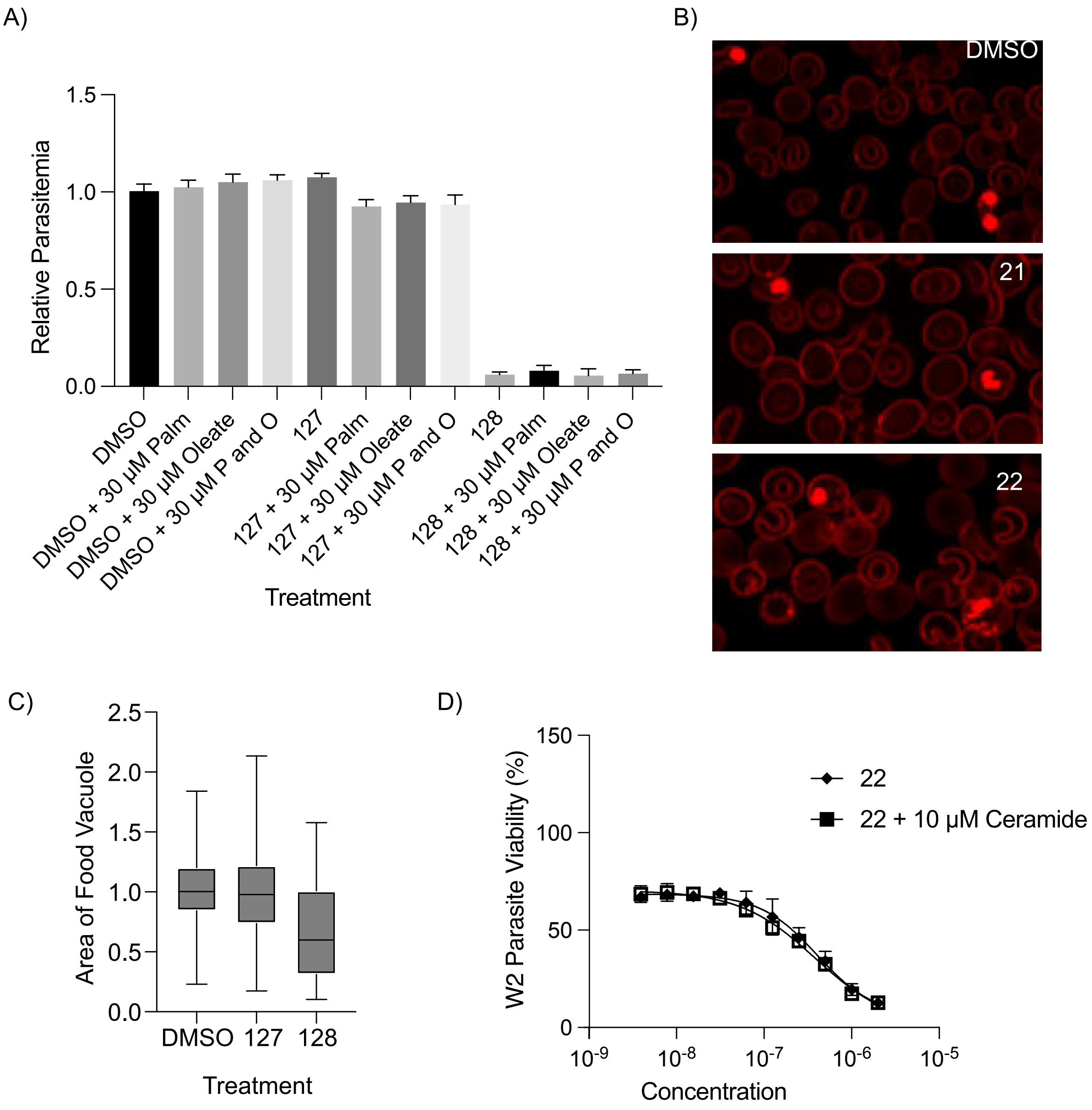


**Figure 6S (Related to Figure 5):** Supplementation does not rescue parasite killing.
(A) Synchronized ring stage parasites were treated with 5 µM of compound **22** as well as with 30 µM of palmitate, oleate, or a combination of the two and grown for 72 hours.
(B) Visualization of food vacuole of Parasites treated with DMSO, compound **21**, or **22** for 16 hours.
(C) Quantification of parasite food vacuole size under the various treatments (N, n=2, 20).
(D) Dose-response curve of parasites treated with either compound **22** or compound **22** with 10 µM ceramide mix.

**Table S1 (Related to Figure 4): Table S1 Primers for the confirmation of P. falciparum conditional mutant lines and WT.**

| Primer Name | Sequence 5'-3' |
|---|---|
| P1 | AGAGGTTCGACGGAAGTATCTGTTATAATG |
| P2 | ACAACTCCAGTGAAAAGTTCTTCTCCT |
| P3 | ATCATTCTTTCCATGCATAATAAATAAAGG |
| P8408 | GTGAGTACATAAATATATTATATAAACTAGACTAGGTTAACTGGCCAAGATCTCCCGGG |
| P8470 | TCCTCCTCAGAAATTAGCTTCTGCTC |
| P8510 | GTTGAGTTGGGAAAATACTGGTGAACTAG |
| P8977 | CGGGTTTAGGGGACGCATGG |
| P8978 | GCTTTTATCACTCAAGCGAAAAGCATGAA |
| P9141 | AGTTAAAATAAGGCTAGTCCGTTATCAACTTG |
| Guide | taatacgactcactataggGAGCTATTTTATCATAAATTgttttagagctagaaatag |

**Table S2 (Related to Figure 6): WGS metric for samples sequenced on MiSeq.**

| | | Drug treated Dd2-PolD | | | | | | Parent |
|---|---|---|---|---|---|---|---|---|
| Sample names | | FL1_B11 | FL1_B6 | FL1_C8 | FL2_A5 | FL2_E7 | FL2_F8 | Dd2_PolD |
| Total reads | | 4,209,799 | 4,503,417 | 3,554,875 | 3,957,932 | 4,217,477 | 4,220,167 | 4,271,890 |
| # Mapped reads | | 3,906,373 | 4,194,721 | 3,303,206 | 3,695,516 | 3,934,380 | 3,925,665 | 3,963,861 |
| Duplication rate | | 28.86% | 34.93% | 28.43% | 28.58% | 27.88% | 27.81% | 28.71% |
| General error rate | | 1.80% | 1.78% | 1.84% | 1.77% | 1.84% | 1.83% | 1.89% |
| Mean mapping quality (Phred) | | 56.57 | 56.64 | 56.61 | 56.62 | 56.59 | 56.57 | 56.54 |
| Depth of coverage | mean | 36.12 | 36.23 | 30.14 | 33.87 | 36.29 | 36.16 | 36.10 |
| | SD | 33.27 | 32.83 | 28.22 | 31.87 | 32.43 | 33.94 | 30.49 |
| % of PF genome with > x no. reads | 1X | 96.03% | 96.08% | 95.89% | 95.97% | 95.99% | 96.04% | 96.07% |
| | 5X | 94.35% | 94.39% | 93.94% | 93.97% | 94.19% | 94.22% | 94.29% |
| | 10X | 92.45% | 92.69% | 91.37% | 91.15% | 92.08% | 92.00% | 92.43% |
| | 30X | 67.10% | 68.23% | 51.66% | 60.16% | 66.17% | 65.84% | 68.06% |

**Table S3 (Related to Figure 6): *P. falciparum* Dd2-B2 asexual blood-stage MIR summary for compound 22.**

| Drug | Dd2-B2 inoculum | Day of Recrudescence | Flask |
|---|---|---|---|
| Compound **22** | $3.3x10^8$ | NA | 1 |
| Compound **22** | $3.3x10^8$ | 23 | 2 |
| Compound **22** | $3.3x10^8$ | NA | 3 |

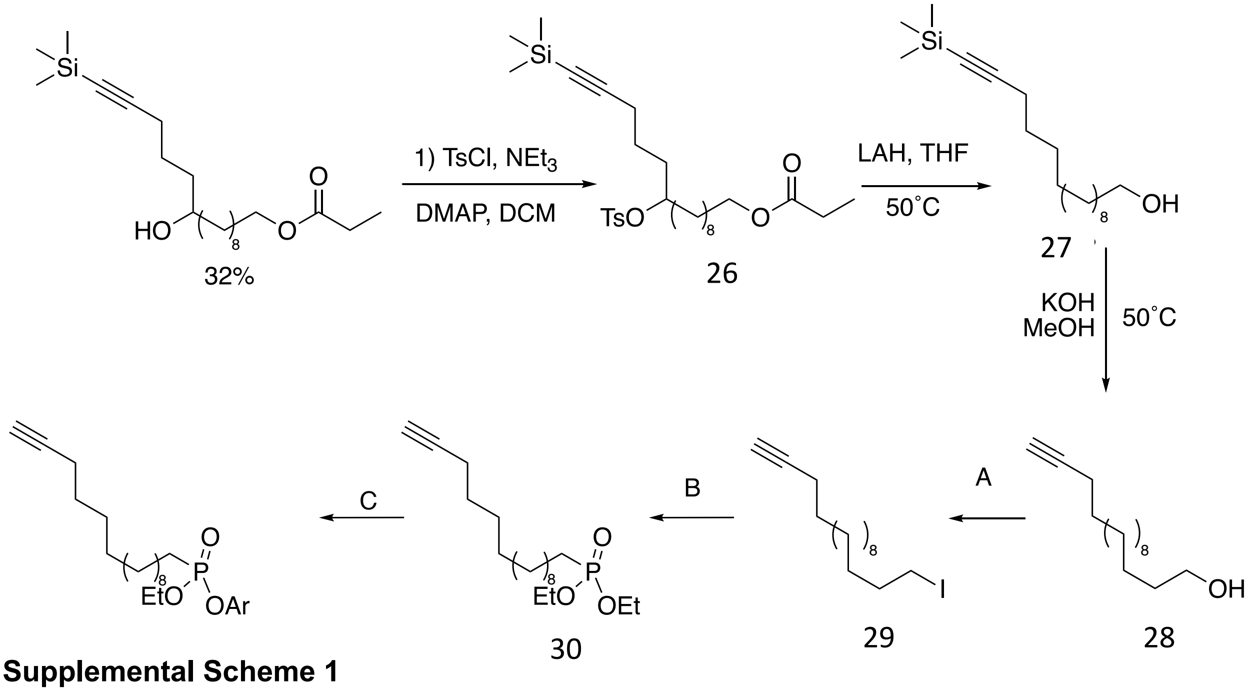


**Supplemental Scheme 1**

NMR spectra were recorded on a Varian 400 MHz (400/100), Varian 500 MHz (500/125) or a Varian Inova 600 MHz (600/150 MHz) equipped with a pulsed field gradient accessory. Chemical shifts are given in ppm (δ) relative to tetramethylsilane as an internal standard. Coupling constants are given in Hz.

3,5-difluorophenyl ethyl hex-5-yn-1-ylphosphonate (1): $^{1}$H NMR (400 MHz, cdcl$_3$) δ 6.83 – 6.76 (m, 2H), 6.62 (dd, *J* = 10.1, 7.8 Hz, 1H), 4.28 – 4.08 (m, 2H), 2.22 (td, *J* = 6.9, 2.6 Hz, 2H), 1.97 – 1.72 (m, 6H), 1.63 (p, *J* = 7.0 Hz, 2H), 1.31 (t, *J* = 7.1 Hz, 3H). $^{13}$C NMR (101 MHz, cdcl$_3$) δ 100.65, 83.39, 68.95, 62.86, 62.79, 28.94, 28.77, 26.04, 24.64, 21.33, 21.27, 17.88, 16.30. $^{31}$P NMR (162 MHz, cdcl$_3$) δ 29.47.

ethyl (4-(fluorosulfonyl)phenyl) hex-5-yn-1-ylphosphonate (2): $^{1}$H NMR (400 MHz, cdcl$_3$) δ 8.03 – 7.94 (m, 2H), 7.52 – 7.41 (m, 2H), 4.31 – 4.07 (m, 2H), 2.22 (td, *J* = 6.9, 2.7 Hz, 2H), 2.06 – 1.91 (m, 2H), 1.96 – 1.74 (m, 2H), 1.70 – 1.58 (m, 2H), 1.31 (t, *J* = 7.1 Hz, 3H). $^{13}$C NMR (101 MHz, cdcl$_3$) δ 130.77, 121.59, 121.54, 116.35, 69.03, 63.14, 63.07, 28.88, 26.26, 24.86, 21.28, 17.88, 16.36, 16.31. $^{31}$P NMR (162 MHz, cdcl$_3$) δ 29.91. $^{19}$F NMR (376 MHz, cdcl$_3$) δ -63.03, -65.87.

2,6-difluorophenyl ethyl hex-5-yn-1-ylphosphonate (3): $^{1}$H NMR (400 MHz, cdcl$_3$) δ 7.08 – 7.01 (m, 1H), 6.97 – 6.89 (m, 2H), 4.37 – 4.18 (m, 2H), 2.22 (td, *J* = 7.0, 2.7 Hz, 2H), 2.09 – 1.96 (m, 2H), 1.94 (t, *J* = 2.7 Hz, 1H), 1.90 – 1.79 (m, 2H), 1.71 – 1.61 (m, 2H), 1.35 (t, *J* = 7.1 Hz, 3H). $^{13}$C NMR (101 MHz, cdcl$_3$) δ 124.84, 112.33, 112.11, 83.62, 68.77, 62.34, 29.15, 28.97, 26.40, 24.99, 21.59, 21.54, 17.98, 17.96, 16.25. $^{31}$P NMR (162 MHz, cdcl$_3$) δ 31.07.

2,6-dimethylphenyl ethyl hex-5-yn-1-ylphosphonate (4): $^{1}$H NMR (400 MHz, cdcl$_3$) δ 7.02 – 6.89 (m, 4H), 4.13 – 4.02 (m, 1H), 3.99 – 3.88 (m, 1H), 2.23 (td, *J* = 7.0, 2.5 Hz, 2H), 1.99 (d, *J* = 7.4

Hz, 1H), 1.98 – 1.82 (m, 3H), 1.66 (p, $J$ = 7.2 Hz, 2H), 1.15 (td, $J$ = 7.1, 1.6 Hz, 4H). $^{13}$C NMR (101 MHz, cdcl$_3$) δ 130.44, 128.94, 128.92, 124.80, 83.62, 68.82, 62.74, 53.44, 29.23, 26.77, 25.34, 21.73, 17.99, 17.46, 16.33. $^{31}$P NMR (162 MHz, cdcl$_3$) δ 28.60.

ethyl (4-(methylthio)phenyl) hex-5-yn-1-ylphosphonate (5): $^1$H NMR (400 MHz, cdcl$_3$) δ 7.21 – 7.17 (m, 1H), 7.12 – 7.08 (m, 2H), 4.22 – 4.05 (m, 2H), 2.42 (d, $J$ = 1.8 Hz, 3H), 2.22 – 2.14 (m, 2H), 1.95 – 1.80 (m, 2H), 1.61 (q, $J$ = 7.2 Hz, 2H), 1.26 (td, $J$ = 7.0, 1.8 Hz, 3H). $^{13}$C NMR (101 MHz, cdcl$_3$) δ 148.38, 134.43, 128.47, 121.05, 83.56, 68.86, 62.51, 29.06, 28.89, 25.89, 24.49, 21.46, 17.93, 16.63, 16.40. $^{31}$P NMR (162 MHz, cdcl$_3$) δ 29.18.

4-(tert-butyl)phenyl ethyl hex-5-yn-1-ylphosphonate (6): $^1$H NMR (400 MHz, cdcl$_3$) δ 7.34 – 7.26 (m, 2H), 7.12 – 7.05 (m, 2H), 4.25 – 4.05 (m, 2H), 2.19 (td, $J$ = 7.0, 2.7 Hz, 2H), 1.94 – 1.71 (m, 6H), 1.67 – 1.57 (m, 2H), 1.27 (s, 9H). $^{13}$C NMR (101 MHz, cdcl$_3$) δ 147.65, 126.55, 119.83, 83.62, 68.80, 62.31, 34.33, 31.38, 29.11, 28.94, 25.87, 24.47, 21.52, 21.47, 17.95, 16.39, 16.33. $^{31}$P NMR (162 MHz, cdcl$_3$) δ 28.91.

ethyl naphthalen-2-yl hex-5-yn-1-ylphosphonate (7): $^1$H NMR (400 MHz, cdcl$_3$) δ 7.37 (t, $J$ = 8.8 Hz, 3H), 7.08 – 6.88 (m, 2H), 3.88 – 3.68 (m, 2H), 1.79 (td, $J$ = 7.0, 4.0 Hz, 2H), 1.59 – 1.37 (m, 5H), 1.26 – 1.20 (m, 1H), 0.91 – 0.86 (m, 3H). $^{13}$C NMR (101 MHz, cdcl$_3$) δ 148.24, 148.16, 133.92, 130.81, 129.84, 127.68, 127.46, 126.67, 125.37, 120.59, 116.83, 83.59, 68.88, 62.58, 29.09, 28.92, 25.99, 24.58, 21.48, 17.96, 16.44 $^{31}$P NMR (162 MHz, cdcl$_3$) δ 29.18.

4-bromo-2,6-dimethylphenyl ethyl hex-5-yn-1-ylphosphonate (8): $^1$H NMR (400 MHz, cdcl$_3$) δ 7.16 – 7.11 (m, 2H), 4.14 – 4.01 (m, 1H), 4.01 – 3.87 (m, 1H), 2.31 (q, $J$ = 0.8 Hz, 9H), 2.23 (td, $J$ = 6.9, 2.6 Hz, 2H), 2.02 – 1.78 (m, 4H), 1.72 – 1.60 (m, 2H), 1.17 (t, $J$ = 7.1 Hz, 3H). $^{13}$C NMR (101 MHz, cdcl$_3$) δ 146.85, , 132.75, 131.53, 117.57, 83.55, 68.89, 62.98, 29.18, 26.79, 25.36, 21.67, 17.99, 17.35, 16.39. $^{31}$P NMR (162 MHz, cdcl$_3$) δ 29.04.

ethyl (perfluorophenyl) hex-5-yn-1-ylphosphonate (9): $^1$H NMR (400 MHz, cdcl$_3$) δ 4.36 – 4.17 (m, 2H), 2.23 (td, $J$ = 7.0, 2.8 Hz, 2H), 2.12 – 1.77 (m, 6H), 1.65 (p, $J$ = 7.0 Hz, 2H), 1.36 (t, $J$ = 7.0 Hz, 3H). $^{13}$C NMR (101 MHz, cdcl$_3$) δ 81.06, 66.42, 66.15, 58.63, 58.56, 26.54, 26.37, 26.19, 23.46, 22.35, 22.03, 18.77, 15.40, 13.73. $^{31}$P NMR (162 MHz, cdcl$_3$) δ 32.67.

[1,1'-biphenyl]-4-yl ethyl hex-5-yn-1-ylphosphonate (10): $^1$H NMR (400 MHz, cdcl$_3$) δ 7.57 – 7.48 (m, 4H), 7.46 – 7.36 (m, 2H), 7.36 – 7.30 (m, 1H), 7.30 – 7.22 (m, 2H), 4.30 – 4.05 (m, 2H), 2.22 (tdd, $J$ = 7.0, 2.7, 0.8 Hz, 2H), 1.99 – 1.77 (m, 4H), 1.71 – 1.59 (m, 2H), 1.31 (t, $J$ = 6.9 Hz, 3H). $^{13}$C NMR (101 MHz, cdcl$_3$) δ 150.17, 140.33, 138.08, 128.90, 128.52, 127.39, 127.09, 120.89, 120.85, 83.71, 68.99, 62.63, 29.21, 26.11, 24.70, 21.64, 18.09, 16.49. $^{31}$P NMR (162 MHz, cdcl$_3$) δ 26.55.

ethyl phenyl hex-5-yn-1-ylphosphonate (11): $^1$H NMR (400 MHz, cdcl$_3$) δ 7.34 – 7.27 (m, 2H), 7.22 – 7.10 (m, 3H), 4.25 – 4.05 (m, 2H), 2.20 (tdd, $J$ = 6.9, 2.7, 0.8 Hz, 2H), 1.95 – 1.74 (m, 5H), 1.67 – 1.58 (m, 2H), 1.28 (td, $J$ = 7.1, 0.8 Hz, 3H). $^{13}$C NMR (101 MHz, cdcl$_3$) δ 127.15, 122.26, 117.93, 81.02, 66.25, 59.87, 26.52, 23.39, 21.98, 18.9315.38, 13.82. $^{31}$P NMR (162 MHz, cdcl$_3$) δ 26.31.

4-cyclopentylphenyl ethyl hex-5-yn-1-ylphosphonate (12): $^1$H NMR (400 MHz, cdcl$_3$) δ 7.19 – 7.13 (m, 2H), 7.07 (dt, $J$ = 8.7, 1.2 Hz, 2H), 4.27 – 4.03 (m, 2H), 2.93 (ddd, $J$ = 17.2, 9.7, 7.4 Hz, 1H), 2.20 (tdd, $J$ = 6.9, 2.7, 0.9 Hz, 2H), 2.06 – 1.96 (m, 2H), 1.96 – 1.82 (m, 3H), 1.64 (ddt, $J$ = 14.3, 9.4, 5.2 Hz, 4H), 1.54 – 1.44 (m, 1H), 1.28 (td, $J$ = 7.1, 0.9 Hz, 3H). $^{13}$C NMR (101 MHz, cdcl$_3$) δ 148.40, 143.03, 128.22, 120.13, 120.08, 83.64, 68.81, 62.33, 62.26, 45.24, 34.62,

29.12, 28.95, 25.86, 25.39, 24.46, 21.53, 21.48, 17.97, 16.41, 16.35. $^{31}$P NMR (162 MHz, cdcl$_3$) δ 28.92.

ethyl (4-nitrophenyl) hex-5-yn-1-ylphosphonate (13): $^{1}$H NMR (400 MHz, cdcl$_3$) δ 8.27 – 8.20 (m, 2H), 7.41 (m, 2H), 4.32 – 4.09 (m, 2H), 2.24 (td, *J* = 6.9, 2.6 Hz, 2H), 2.03 – 1.97 (m, 2H), 1.97 – 1.76 (m, 4H), 1.67 (q, *J* = 7.0 Hz, 4H), 1.33 (t, *J* = 7.1 Hz, 3H). $^{31}$P NMR (162 MHz, cdcl$_3$) δ 27.14.

4-cyanophenyl ethyl hex-5-yn-1-ylphosphonate (14): $^{1}$H NMR (400 MHz, cdcl$_3$) δ 7.65 – 7.57 (m, 3H), 7.35 – 7.27 (m, 3H), 4.29 – 4.02 (m, 4H), 2.20 (td, *J* = 6.9, 2.6 Hz, 3H), 1.96 – 1.92 (m, 1H), 1.92 – 1.71 (m, 7H), 1.67 – 1.53 (m, 4H), 1.26 (s, 3H). $^{13}$C NMR (101 MHz, cdcl$_3$) δ 154.04, 134.08, 121.42, 121.38, 118.27, 108.61, 83.39, 68.99, 62.93, 62.86, 28.93, 28.76, 26.18, 24.78, 21.34, 21.29, 17.91, 17.89, 16.39, 16.33. $^{31}$P NMR (162 MHz, cdcl$_3$) δ 29.60.

ethyl (4-methoxyphenyl) hex-5-yn-1-ylphosphonate (15): $^{1}$H NMR (400 MHz, cdcl$_3$) δ 7.12 – 7.04 (m, 3H), 6.84 – 6.75 (m, 3H), 4.18 – 4.07 (m, 2H), 3.74 (s, 3H), 2.18 (td, *J* = 7.0, 2.7 Hz, 3H), 1.88 – 1.75 (m, 6H), 1.61 (q, *J* = 7.5 Hz, 4H), 1.26 (d, *J* = 7.1 Hz, 3H). $^{13}$C NMR (101 MHz, cdcl$_3$) δ 156.54, 121.37, 121.33, 114.63, 114.62, 83.62, 68.81, 62.41, 62.34, 55.58, 29.11, 28.94, 25.78, 24.38, 21.51, 21.46, 17.95, 17.94, 16.42, 16.36. $^{31}$P NMR (162 MHz, cdcl$_3$) δ 29.20.

ethyl (4-isopropoxyphenyl) hex-5-yn-1-ylphosphonate (16): $^{1}$H NMR (400 MHz, cdcl$_3$) δ 7.17 – 7.10 (m, 2H), 7.10 – 7.03 (m, 2H), 4.25 – 4.03 (m, 2H), 2.19 (td, *J* = 7.0, 2.8 Hz, 2H), 1.80 (s, 1H), 1.61 (m, *J* = 7.3 Hz, 2H), 1.30 – 1.17 (m, 9H), 0.88 – 0.77 (m, 2H). $^{13}$C NMR (101 MHz, cdcl$_3$) δ 127.56, 120.20, 120.16, 83.63, 68.80, 62.33, 62.26, 33.47, 29.11, 28.94, 25.87, 24.46, 24.04, 21.52, 21.47, 17.96, 17.94, 16.39, 16.33. $^{31}$P NMR (162 MHz, cdcl$_3$) δ 28.92.

ethyl (4-(pentafluoro-sulfaneyl)phenyl) hex-5-yn-1-ylphosphonate (17): $^{1}$H NMR (400 MHz, cdcl$_3$) δ 7.76 – 7.65 (m, 2H), 7.33 – 7.26 (m, 2H), 4.30 – 4.06 (m, 2H), 2.22 (td, *J* = 6.9, 2.6 Hz, 2H), 1.99 – 1.73 (m, 5H), 1.70 – 1.58 (m, 2H), 1.31 (t, *J* = 7.1 Hz, 3H). $^{13}$C NMR (101 MHz, cdcl$_3$) δ 127.86, 120.49, 120.44, 83.40, 68.96, 62.85, 62.78, 28.96, 28.78, 26.12, 24.72, 21.36, 21.31, 17.91, 16.39, 16.33. $^{31}$P NMR (162 MHz, cdcl$_3$) δ 29.55.

4-acetylphenyl ethyl hex-5-yn-1-ylphosphonate (18): $^{1}$H NMR (400 MHz, cdcl$_3$) δ 7.96 – 7.89 (m, 2H), 7.31 – 7.23 (m, 2H), 4.27 – 4.04 (m, 2H), 2.55 (d, *J* = 0.4 Hz, 3H), 2.20 (td, *J* = 7.0, 2.7 Hz, 2H), 1.98 – 1.89 (m, 2H), 1.80 (s, 2H), 1.69 – 1.56 (m, 2H), 1.28 (t, *J* = 7.1 Hz, 3H). $^{13}$C NMR (101 MHz, cdcl$_3$) δ 196.80, 154.42, 133.77, 130.39, 120.45, 120.41, 83.47, 68.93, 62.76, 62.68, 29.01, 28.83, 26.57, 26.14, 24.74, 21.41, 21.36, 17.93, 17.92, 16.40, 16.34. $^{31}$P NMR (162 MHz, cdcl$_3$) δ 29.20.

ethyl mesityl hex-5-yn-1-ylphosphonate (19): $^{1}$H NMR (400 MHz, cdcl$_3$) δ 6.86 – 6.74 (m, 2H), 4.14 – 4.03 (m, 1H), 3.93 (dddd, *J* = 14.9, 10.2, 7.5, 5.1 Hz, 1H), 2.29 (s, 9H), 2.25 – 2.18 (m, 7H), 2.02 – 1.77 (m, 8H), 1.76 – 1.59 (m, 3H), 1.17 (ddd, *J* = 7.2, 6.3, 1.0 Hz, 3H). $^{13}$C NMR (101 MHz, cdcl$_3$) δ 129.97, 129.54, 129.52, 83.68, 68.80, 62.67, 29.27, 29.09, 26.75, 25.32, 21.76, 21.71, 20.59, 18.02, 18.01, 17.40, 16.39, 16.33. $^{31}$P NMR (162 MHz, cdcl$_3$) δ 28.67.

ethyl (2,3,5,6-tetrafluorophenyl) hex-5-yn-1-ylphosphonate (20): $^{1}$H NMR (400 MHz, cdcl$_3$) δ 6.90 (ttd, *J* = 10.0, 7.2, 1.2 Hz, 1H), 4.41 – 4.19 (m, 2H), 2.24 (td, *J* = 6.9, 2.7 Hz, 2H), 2.12 – 1.99 (m, 2H), 1.99 – 1.79 (m, 3H), 1.73 – 1.61 (m, 2H), 1.38 (t, *J* = 7.1 Hz, 3H).

2,6-difluorophenyl ethyl pentadecylphosphonate (21): $^{1}$H NMR (400 MHz, cdcl$_3$) δ 7.10 – 7.00 (m, 1H), 6.93 (t, *J* = 8.0 Hz, 2H), 4.40 – 4.18 (m, 2H), 2.06 – 1.94 (m, 2H), 1.69 (dd, *J* = 16.6, 7.7 Hz, 4H), 1.29 (d, *J* = 43.2 Hz, 26H), 0.86 (t, *J* = 6.7 Hz, 3H). $^{31}$P NMR (162 MHz, cdcl$_3$) δ 31.97.

3,5-difluorophenyl ethyl pentadecylphosphonate (22): $^{1}$H NMR (400 MHz, cdcl$_3$) δ 6.79 (ddd, *J* = 7.8, 2.4, 1.2 Hz, 2H), 6.61 (t, *J* = 8.9 Hz, 1H), 4.28 – 4.07 (m, 2H), 1.87 (dt, *J* = 18.0, 8.3 Hz, 2H), 1.64 (s, 3H), 1.36 – 1.22 (m, 26H), 0.86 (t, *J* = 6.3 Hz, 3H). $^{31}$P NMR (162 MHz, cdcl$_3$) δ 30.32. $^{19}$F NMR (376 MHz, cdcl$_3$) δ -107.84 (t, *J* = 8.0 Hz), -110.00 (t, *J* = 7.9 Hz ).

ethyl phenyl pentadecylphosphonate (23): $^{1}$H NMR (400 MHz, cdcl$_3$) δ 7.40 – 7.24 (m, 3H), 7.18 (s, 2H), 7.23 – 7.09 (m, 3H), 4.26 – 4.07 (m, 2H), 1.94 – 1.81 (m, 2H), 1.73 – 1.62 (m, 2H), 1.27 (d, *J* = 16.7 Hz, 28H), 0.88 (t, *J* = 6.6 Hz, 3H). $^{31}$P NMR (162 MHz, cdcl$_3$) δ 27.11.

1-iodopentadecane (24): $^{1}$H NMR (400 MHz, cdcl$_3$) δ 3.17 (t, *J* = 7.1 Hz, 2H), 1.80 (p, *J* = 7.1 Hz, 2H), 1.40 – 1.34 (m, 2H), 1.25 (d, *J* = 3.9 Hz, 22H), 0.86 (t, *J* = 6.7 Hz, 3H). $^{13}$C NMR (101 MHz, cdcl$_3$) δ 33.56, 31.90, 30.49, 29.66, 29.65, 29.63, 29.59, 29.52, 29.40, 29.33, 28.52, 22.67, 14.09, 7.33.

diethyl pentadecylphosphonate (25): $^{1}$H NMR (400 MHz, cdcl$_3$) δ 4.34 – 3.85 (m, 1H), 1.79 – 1.63 (m, 1H), 1.54 – 0.97 (m, 12H), 0.86 (t, *J* = 6.6 Hz, 1H).

10-(tosyloxy)-15-(trimethylsilyl)pentadec-14-yn-1-yl propionate (26): $^{1}$H NMR (400 MHz, cdcl$_3$) δ 7.78 (dd, *J* = 8.3, 3.3 Hz, 2H), 7.32 (t, *J* = 8.1 Hz, 2H), 4.08 (dt, *J* = 29.3, 6.4 Hz, 2H), 2.43 (d, *J* = 2.1 Hz, 3H), 2.35 – 2.21 (m, 2H), 2.13 (t, *J* = 6.9 Hz, 1H), 1.83 (p, *J* = 6.6 Hz, 1H), 1.72 – 1.37 (m, 4H), 1.37 – 1.08 (m, 6H), 0.14 – 0.04 (m, 9H). $^{13}$C NMR (101 MHz, cdcl$_3$) δ 129.83, 129.65, 127.88, 127.67, 68.88, 27.86, 25.87, 21.61, 16.06, 0.03.

15-(trimethylsilyl)pentadec-14-yn-1-ol (27): $^{1}$H NMR (400 MHz, cdcl$_3$) δ 3.62 (t, *J* = 6.6 Hz, 2H), 2.19 (t, *J* = 7.2 Hz, 2H), 1.58 – 1.45 (m, 4H), 1.25 (s, 7H), 0.13 (s, 9H). $^{13}$C NMR (101 MHz, cdcl$_3$) δ 63.07, 32.79, 29.59, 29.58, 29.56, 29.55, 29.45, 29.40, 29.05, 28.77, 28.61, 25.71, 19.83.

pentadec-14-yn-1-ol (28): $^{1}$H NMR (400 MHz, cdcl$_3$) δ 3.62 (t, *J* = 6.6 Hz, 2H), 2.16 (td, *J* = 7.1, 2.7 Hz, 2H), 1.60 – 1.44 (m, 4H), 1.37 (s, 3H), 1.25 (d, *J* = 3.9 Hz, 17H). $^{13}$C NMR (101 MHz, cdcl$_3$) δ 67.99, 63.07, 32.78, 29.57, 29.46, 29.39, 29.08, 28.73, 28.47, 25.70, 18.37.

15-iodopentadec-1-yne (29): $^{1}$H NMR (400 MHz, cdcl$_3$) δ 4.11 – 4.02 (m, 4H), 2.16 (td, *J* = 7.1, 2.7 Hz, 2H), 1.77 – 1.64 (m, 2H), 1.64 – 1.44 (m, 4H), 1.40 – 1.24 (m, 19H). $^{13}$C NMR (101 MHz, cdcl$_3$) δ 67.99, 30.67, 29.55, 29.46, 29.35, 29.07, 28.73, 28.46, 24.96, 18.37, 16.41. $^{31}$P NMR (162 MHz, cdcl$_3$) δ 32.69.

2,6-difluorophenyl ethyl pentadec-14-yn-1-ylphosphonate (21-probe): $^{1}$H NMR (400 MHz, cdcl$_3$) $^{1}$H NMR (400 MHz, cdcl$_3$) δ 7.11 – 6.99 (m, 1H), 6.98 – 6.88 (m, 2H), 4.37 – 4.17 (m, 2H), 2.16 (td, *J* = 7.1, 2.6 Hz, 2H), 2.06 – 1.89 (m, 3H), 1.71 (q, *J* = 13.7 Hz, 2H), 1.58 (s, 1H), 1.49 (q, *J* = 7.2 Hz, 2H), 1.36 (dt, *J* = 14.1, 7.4 Hz, 9H), 1.28 – 1.22 (m, 15H). $^{13}$C NMR (101 MHz, cdcl$_3$) δ 164.28, 161.80, 104.72, 104.44, 100.79, 100.27, 84.76, 67.99, 62.69, 31.55, 30.48, 30.31, 29.53, 29.44, 29.28, 29.06, 28.98, 28.72, 28.45, 26.50, 25.11, 22.61, 22.18, 18.36, 16.36, 14.07. $^{31}$P NMR (162 MHz, cdcl$_3$) δ 31.86 (t, *J* = 1.9 Hz).

3,5-difluorophenyl ethyl pentadec-14-yn-1-ylphosphonate (22-probe): $^{1}H$ NMR (400 MHz, $cdcl_3$) δ 6.81 – 6.75 (m, 2H), 6.61 (tt, *J* = 8.9, 2.3 Hz, 1H), 4.27 – 4.06 (m, 2H), 2.15 (td, *J* = 7.1, 2.6 Hz, 2H), 1.93 – 1.80 (m, 3H), 1.64 (ddt, *J* = 15.6, 11.9, 7.5 Hz, 2H), 1.55 – 1.44 (m, 2H), 1.37 (dq, *J* = 12.5, 7.1 Hz, 4H), 1.33 – 1.19 (m, 22H). $^{13}C$ NMR (101 MHz, $cdcl_3$) δ 164.28, 161.80, 104.72, 104.44, 100.79, 100.53, 100.27, 84.76, 67.99, 62.69, 31.55, 30.48, 30.31, 29.53, 29.44, 29.28, 29.06, 28.98, 28.72, 28.45, 26.50, 25.11, 22.61, 22.18, 18.36, 16.36, 14.07. $^{31}P$ NMR (162 MHz, $cdcl_3$) δ 30.29.
